# An allelic series of spontaneous mutations in *Rorb* cause a gait phenotype, retinal abnormalities, and transcriptomic changes relevant to human neurodevelopmental conditions

**DOI:** 10.1101/2021.11.23.468991

**Authors:** George C. Murray, Jason Bubier, Oraya J. Zinder, Belinda Harris, James Clark, Mia-Cara Christopher, Courtany Hanley, Harianto Tjong, Meihong Li, Chew Yee Ngan, Laura Reinholdt, Robert W. Burgess, Abby L.D. Tadenev

## Abstract

*Rorb* encodes the Retinoic Acid Receptor-related orphan receptor beta. Mutations in either of the two transcripts of *Rorb* cause defects in multiple systems, including abnormal photoreceptor abundance and morphology in the retina and a characteristic “high-stepper” or “duck-like” gait arising from dysfunction of interneurons in the spinal cord. *Rorb* is also important for cortical development and cell fate specification in mice. *Rorb* variants segregate with epilepsy and comorbidities such as intellectual disability in numerous clinical cases. Here we describe five mouse strains with spontaneous mutations in *Rorb* identified by their gait phenotype. These mutations affect different domains and isoforms of *Rorb*, which correspond to the spectrum of anatomical and physiological phenotypes exhibited by these mice. Gene set analysis in *Rorb* mutants implicates pathways associated with development and nervous system function, and differential gene expression analysis indicates changes in numerous genes related to epilepsy, bipolar disorder, and autism spectrum disorder (ASD). Many of these genes and their protein products are known to interact during synapse formation and neuronal activity. These findings further illuminate the role of *Rorb* in nervous system development, provide further evidence for an association between *Rorb* and several neurological conditions, and describe an allelic series of *Rorb* mutant mice that will be useful for dissecting thalamocortical afferent (TCA) development, neural cell fate determination, and as animal models exhibiting transcriptomic shifts in neurological conditions such as epilepsy, bipolar disorder, and ASD.

■ Five mutant mice with a characteristic high-stepper gait phenotype have mutations in *Rorb*
■ These allelic series mutations manifest in a spectrum of anatomical and physiological abnormalities
■ Gene expression data suggest involvement of pathways related to neurodevelopmental disorders

## INTRODUCTION

Retinoic Acid Receptor-related orphan receptor beta *(RORB)* encodes a transcription factor located on human chromosome 9q22, a region syntenic with mouse chromosome 19. It is widely expressed in the nervous system with particularly high levels in the cerebral cortex and retina of humans and mice(Andre *et al*. 1998a; Lindskog 2015; Uhlen *et al*. 2015). Periodic fluctuations in the expression of *Rorb* have been detected in regions with circadian involvement, such as the suprachiasmatic nucleus, the retina, and the pineal gland (Andre *et al*. 1998a). In mice and humans, *Rorb* has two isoforms with alternative first exons, and in mice each exhibits a degree of tissue-specificity and temporally divergent expression during development(Andre *et al*. 1998b; Liu *et al*. 2013a). In mice, *Rorb* knockout causes a characteristic gait phenotype referred to as “high stepper” or “duck gait,” which was associated with electrophysiological dysfunction in a subpopulation of *Pax2+* inhibitory spinal interneurons that require RORB for proper gating of input and rhythmic output from the central pattern generator of the spinal cord(Koch *et al*. 2017). A mechanism whereby splice-site mutations in *RORB* cause reduction of RORB-positive neurons in the spinal cord and abnormal spinal interneuron differentiation was recently shown to underlie abnormal gait in *sauteur d’Alfort* rabbits(Carneiro *et al*. 2021). Delayed onset of fertility in male mice with *Rorb* mutations has also been described previously(Andre *et al*. 1998b). Mice lacking *Rorb* also display a pronounced lack of rod photoreceptors and abnormal cone photoreceptors without outer segments, leading to disorganization of the retina and suggesting a role for RORB in cell fate determination(Jia *et al*. 2009). These retinal phenotypes are likely related to *Rorb* induction of *neural retina leucine zipper factor (Nrl)* expression, which plays a critical role in the cone:rod cell fate choice(Fu *et al*. 2014). *Rorb* expression is also required for induction of opsin in cone photoreceptors(Srinivas *et al*. 2006). Further involvement of *Rorb* in the development of sensory systems of sensory signaling is evidenced by its embryonic requirement for proper formation of whisker barrels in the somatosensory cortex of mice(Jabaudon *et al*. 2012) and its role in establishment of long-range axonal projections from the superior colliculus to the dorsal lateral geniculate nucleus and the pretectum(Byun *et al*. 2019). Disruption of cytoarchitectural patterning due to loss of *Rorb* may occur via a shift in cellular identity initiated through the downstream RORB target *Thsd7a* and/or changes in chromatin accessibility during development(Clark *et al*. 2020).

The two *Rorb* isoforms, referred to as *Rorb1* and *Rorb2*, are expressed via different promoters at different embryonic and postnatal timepoints. In the retina, *Rorb1* mRNA is present throughout mid-embryonic stages and in the first postnatal week, before declining, whereas *Rorb2* is expressed in late embryonic stages and rises during the first postnatal week(Liu *et al*. 2013a). *Rorb1* is thought to be the predominant isoform in the brain, where it is expressed as early as embryonic day 15 and is maintained into adulthood where its expression fluctuates along with components of the molecular pacemaker in regions involved in circadian rhythm(Schaeren-Wiemers *et al*. 1997).

Recently, an association between allelic variation in *RORB* and epilepsy has emerged in human genetic studies. Rudolf and colleagues describe six nonsense variants, one substitution, twelve deletions, and one balanced translocation in *RORB* amongst patients with various forms of epilepsy and epilepsy-related conditions, drawing on both clinical cases and existing data(Gnirke *et al*. 2009; Lesca *et al*. 2013; Rudolf *et al*. 2016). Three additional epileptic patients with documented deletions in 9q21.13 containing *RORB* were also described(Coppola *et al*. 2019). Screening of large patient cohorts revealed 14 individuals with *RORB* variants, 11 of whom have been diagnosed with epilepsy. Notably, two of the 14 individuals were asymptomatic, suggesting that the location of sequence variation within *RORB*, genetic background, and/or environmental factors may influence penetrance(Sadleir *et al*. 2020). Evidence for an association between *RORB* and other neurodevelopmental disorders is also present but less abundant. An exome sequencing study of autism spectrum disorder (ASD), including samples from 11,986 affected individuals, identified an association between ASD and *RORB* variants that passed a FDR < 0.01 threshold(Satterstrom *et al*. 2020). Gait disturbances are frequently observed in ASD, though this relationship is generally thought to involve the cerebellum and basal ganglia(Kindregan *et al*. 2015). Finally, there is evidence for involvement of *RORB* and other circadian-related genes in bipolar disorder(Mansour *et al*. 2009a; McGrath *et al*. 2009; McCarthy *et al*. 2012; Lai *et al*. 2015). The basis for involvement of *RORB* in multiple neurological conditions is unclear but likely involves its developmental role in the central nervous system.

Here we describe an allelic series in mice with mutations in *Rorb* that will be useful for the further examination of the genetic pathways impacted by *Rorb* mutations, as well as clarification of the role of each *Rorb* isoform in the different phenotypic abnormalities observed. These five *Rorb* mutant strains were identified on the basis of their gait phenotype and arose spontaneously at The Jackson Laboratory. Within these alleles, we observed mutations affecting *Rorb1* exclusively, as well as a mutation impacting both isoforms. We examined retinal histology, gait phenotype, tissue-specific expression of *Rorb*, differential gene expression, and pathway/network involvement in several of these mutants and discuss their potential utility in the study of neurological disorders.

## MATERIALS & METHODS

### A. Animals

Spontaneous mutant high-stepping mice were identified at The Jackson Laboratory as part of the Mouse Mutant Resource, which monitors breeding colonies for the appearance of abnormal phenotypes. These phenotypes are then tested for heritability and strains with heritable phenotypes are established as distinct lines. Mice carrying targeted mutations were obtained from The Jackson Laboratory. Official strain designations and the abbreviated names used in this paper are shown in Table 2.

*Rorb^h1^* arose on the C57BL/6J background, while both *Rorb^h2^* and *Rorb^h3^* arose on B6(Cg)-*Tyr*^*c*-2*J*^/J, which is C57BL/6J with the addition of a mutation in the tyrosinase gene (“albino B6”). *Rorb^h4^* arose on B6.129S7-*Il1r1^tm1Imx^*/J, a C57BL/6J congenic strain harboring a mutation in the *Il1r1* gene. *Rorb^h5^* arose on *DBA*/1*J*. Animals were housed under standard conditions at The Jackson Laboratory and all procedures were performed in accordance with The Guide on the Care and Use of Laboratory Animals and were approved by the Institutional Animal Care and Use Committee of the Jackson Laboratory.

In order to map the *hstp1* mutation, affected (and presumed homozygous) mice were crossed to wild-type (WT) BALB/cByJ mice to produce obligate heterozygous F1 mice, which were subsequently intercrossed to produce F2 mice. The *hstp1* allele had previously been mapped to a 15 Mb region of chromosome 19 between the markets D19MIT128 and D19MIT134. 1056 additional F2 mice further narrowed this interval to a 0.7 Mb interval between D19MIT41 and D19MIT113.

### B. Genotyping

Toe biopsies were obtained from mice within the first week of life and were digested overnight in a Proteinase K solution. This solution was used directly for PCR and products were visualized using standard electrophoretic techniques (*Rorb^h1^, Rorb^Cre^, Scnn1a^Cre^*, and *ROSA^tdT^*) or analyzed by Sanger sequencing (*Rorb^h4^, Rorb^h5^*). The primers used were as follows: *Rortb^h1^*: Forward = AGGAGGAGGAATGGGAAGAA, Reverse = TGTGAAGCCCTGCTATCCTT; *Rorb^h4^: Forward* = TCATGTGACAGGGGTCTGAA, Reverse = GCCTTTGCATTGTCCAAAAA; *Rorb^h5^*: Forward = GGTAGTTTACTTGTAACAGGC, Reverse = TTTCCAATCTGGGCAGCAGC; *Rorb^Cre^*: Forward = AACTTGCATGGGGAGAAGC, Reverse (WT allele) = GTTCTCGTCCCCTTCATTTG; Reverse (*Cre* allele) = CCCTCACATTGCCAAAAGAC; *Scnn1a^Cre^* (generic *Cre*): Forward = GCATTACCGGTCGATGCAACGAGTG, Reverse = GAGTGAACGAACCTGGTCGAAATCA; *ROSA^tdT^* (WT allele): Forward = AAGGGAGCTGCAGTGGAGTA, Reverse = CCGAAAATCTGTGGGAAGTC; *ROSA^tdT^* (tdT allele): Forward = GGCATTAAAGCAGCGTATCC, Reverse = CTGTTCCTGTACGGCATGG.

### C. Tissue collection

All tissues were collected following CO_2_ euthanasia.

#### Retina

Eyes were removed, and the lens and cornea were separated from the eye cup using fine scissors. For histology, the entire eye cup was fixed in 4% PFA for 4 hours at 4°C then immersed in 30% sucrose overnight at 4°C. Eye cups were then frozen in OCT and cryosectioned. For qPCR, retinas were gently removed from the eye cup and placed in RNALater (Life Technologies) and incubated overnight at 4°C. Tissue was then removed from the RNALater and frozen at −80°C.

#### Brain

The brain was removed and cut in half sagittally. For histology, one half was then fixed overnight in 4% PFA at 4°C. Tissue was then mounted in an agarose block and sectioned at 100um using a vibratome. For qPCR, the half brain was cut into ~6 pieces, placed in RNALater (Life Technologies), and incubated overnight at 4°C. Tissue was then removed from the RNALater and frozen at −80°C.

#### Spinal cord

The entire spinal column was removed and fixed 24-48h in 4% PFA at 4°C. The spinal cord was then removed from the vertebral column, immersed overnight in 30% sucrose, dissected to isolate the lumbar region, frozen in OCT, and cryosectioned at 20 μm.

#### Femoral nerve

Motor and sensory branches of the femoral nerve were removed and fixed overnight in 2% glutaraldehyde, 2% paraformaldehyde, in 0.1 M cacodylate buffer at 4°C. Both nerve branches were embedded in plastic and 0.5-μm sections were cut and stained with toluidine blue.

### D. Histology, and immunofluorescence

Hematoxylin/eosin staining was performed on retinal sections according to standard techniques. Antibodies used were as follows: 1:200 rabbit anti-S opsin (Millipore AB5407), 1:250 rabbit anti-calcitonin gene-related peptide (CGRP) (Millipore PC205L-100UL), 1:250 rat anti-myelin basin protein (MBP) (Millipore MAB386); secondary antibodies were Alexa Fluor® conjugates from Life Technologies and used at 1:500. FITC-PNA (Sigma L7381) was used at 0.01mg/ml and Isolectin GS-IB_4_, Alexa Fluor® 647 conjugate (Life Technologies I32450) was used at 1:200. Antibodies were applied in phosphate buffered saline (PBS) containing 0.5% TritonX-100 and 3% fetal bovine serum and incubated on sections overnight at 4°C. Fluorophore-conjugated proteins were applied to sections during incubation with secondary antibody; both of these were diluted in PBS. Retinal sections were imaged using a NanoZoomer 2.0HT (Hematoxylin/eosin staining) or a Zeiss Axio Imager (fluorescently-labeled sections). Both vibratome sections and cryosections from brain and spinal cord were imaged using a Leica SP5 confocal. Single confocal slices are shown.

### E. Axon counting and quantification

For axon counting and axon area measurement, images were captured using a Nikon Eclipse E600 microscope with a 40x objective and Nomarski DIC optics. Automated quantification was performed as described in detail previously(Bogdanik *et al*. 2013). Briefly, with the ImageJ software, the Threshold function was used to highlight axoplasm only on whole nerve sections; the Analyze Particle function was then used to count the number of myelinated axons and their cross-sectional areas in each nerve.

### F. Behavioral testing

#### Hot plate testing

Mice were brought into the testing room to acclimate for at least 10 minutes prior to testing. Animals were then individually placed on the hot plate set to 55°C; a Plexiglas cylinder was then placed around the animal to keep them on the plate. The mouse was monitored for a maximum of 30 seconds and the time until first hindpaw lick or flick was recorded. Control mice were a mix of *Rorb^+/+^* and *Rorb^+/h1^*.

#### Von Frey testing

Mice were removed from main housing facility and brought to the testing room to habituate for no less then 30 minutes prior to being placed in testing apparatus. Mice were placed in a plexiglas enclosure (9 cm L x 5 cm W x 5 cm H) on a wire-mesh floor with opaque sides and clear front for observation. Mice were allowed to habituate to the testing chamber for 120 min prior to administration. Von Frey nylon monofilaments purchased from Stoelting, catalogue number 18011 were used to test for mechanical sensitivity thresholds. Each monofilament was calibrated with a scale prior to testing.

A series of eight Von Frey fibers with logarithmically increasing stiffness (0.067 to 9.33 g) were used. Fibers were positioned perpendicular to the plantar surface of the hindpaw, alternating between left and right with a minimum of 5 minutes between responses. Enough pressure was applied to cause a slight bend in the filament and held for 6-8 seconds or until a response is elicited. Tests continued until a maximum of 9 determinations on each paw. Mechanical sensitivity thresholds were determined of each mouse over the course of 3 consecutive days. The threshold force required to elicit a response (median 50% paw withdrawal was determined using the up-down method(Chaplan *et al*. 1994).

Upon completion of the day’s trial, mice were returned to home cage and monitored up to 60 minutes prior to returning to main housing room.

### G. 5’ RACE

5’ RACE was performed following manufacturer’s protocol (Invitrogen 18374-041) on total RNA harvested from WT and *Rorb^h1/h1^* brain. The *Rorb*-specific primer used for first strand synthesis was ACGTGATGACTCGTAGTGGA. For PCR of the RACE product, the primer GCCACAAATTTTGCATGGTA was used with the included abridged anchor primer.

### H. qPCR

Retinal and brain tissue was thawed and homogenized in Trizol using a mechanical homogenizer and total RNA was extracted using the manufacturer’s protocol (Life Technologies 15596). cDNA synthesis (Life Technologies 18080-400) was performed using 1ug total RNA and a 50/50 mix of oligo dT and random hexamers. qPCR was then performed using 1ul of the resulting cDNA in a 20ul reaction using standard SYBR green reagents (Life Technologies 4309155) on a ViiA™ 7 Real-Time PCR System. Results were analyzed using the ΔΔCt method, normalizing to GAPDH expression and to the average of the two WT control groups. Data are presented as arbitrary units, which were calculated as 100 times the fold change. Primers used for qPCR were as follows: *Rorb1/Rorb2* common primer set: Forward = AGGAACCGTTGCCAACACTG, Reverse = GACATCCTCCCGAACTTTACAG; *Rorb1*-specific: Forward = GGCTGGGAGCTTCATGACTA, Reverse = ACGTGATGACTCCGTAGTGGA; *Rorb2*-specific: Forward = CCAGCCCAAAACTAAAGCTG, Reverse = ACGTGATGACTCCGTAGTGGA; *GAPDH*: Forward = AGGTCGGTGTGAACGGATTTG, Reverse = TGTAGACCATGTAGTTGAGGTCA.

### I. RNAseq

Whole brains were isolated, cut into 8-10 pieces, and immersed overnight in RNALater at 4°C, at which point they were transferred to −80°C for storage until being processed. Frozen brain tissues were homogenized in TRIzol (ThermoFisher Scientific) using a gentleMACS dissociator (Miltenyi Biotec Inc). Total RNA was isolated by the Genome Technologies core facility at The Jackson Laboratory using the TRIzol Plus RNA Purification Kit (ThermoFisher Scientific), according to manufacturers’ protocols, including an optional DNase digestion step. Sample concentration and quality were assessed using the Nanodrop 2000 spectrophotometer (Thermo Scientific) and the RNA 6000 Nano LabChip assay (Agilent Technologies). Poly(A) RNA-seq libraries were constructed by the Genome Technologies core facility at The Jackson Laboratory using the TruSeq RNA Library Prep Kit v2 (Illumina). Libraries were checked for quality and concentration using the DNA 1000 LabChip assay (Agilent Technologies) and quantitative PCR (KAPA Biosystems), according to the manufacturers’ instructions. Libraries were pooled and sequenced 100 bp paired-end on the HiSeq 2500 (Illumina) using TruSeq SBS Kit v3-HS (Illumina). RNAseq data are available on GEO using accession number GSE145852.

### J. Mouse neural progenitor cells for ChIA-PET

Mouse embryos were harvested at E12.5 for neural progenitor harvest. Neural progenitors were plated in 6 well plates coated with poly-L-ornithine and grown in N2B27 media for 9 days or until confluent. 10 million cells were dual-crosslinked with 1.5mM EGS (#21565, Thermo Fisher) for 45 min followed by 1% formaldehyde (F8775, Sigma) for 10 min at room temperature and then quenched with 0.2M Glycine (G8898, Sigma) for 5 min. The crosslinked cells were washed with PBS twice and lysed in 100ul 0.55% SDS with incubation at room temperature, 62°C and 37°C sequentially for 10 min each, which was followed by 37°C for 30 min with addition of 50ul 10% Triton-X 100 to quench SDS and 37°C overnight with addition of 40ul AluI (R0137L, NEB) total, 50μl 10× CutSmart buffer to fragmentize the chromatin. The pelleted digested nuclei were resuspended in 500ul A-tailing solution containing 50μl 10× CutSmart buffer, 10μl BSA (B9000S, NEB), 10μl 10mM dATP (N0440S, NEB), 10μl Klenow (3’-5’ exo-) (M0202L, NEB), and 420μl H_2_O with 1 hr incubation at room temperature and then subjected to proximity ligation by adding 200μl 5× ligation buffer (B6058S, NEB), 6μl biotinylated bridge linker (200ng/ul), 10μl T4 DNA ligase (M0202L, NEB) and incubating at 16°C overnight. The ligated chromatins were then sheared by sonication and immunoprecipitated with anti-CTCF (Active Motive #61311). The immunoprecipitated DNA tagmentation, biotin selection, library preparation and sequencing were performed as described(Zhang *et al*. 2013; Tang *et al*. 2015).

### K. ChIA-PET data analysis

ChIA-PET data were processed with ChIA-PET Utilities, a scalable re-implementation of ChIA-PET Tools(Lee *et al*. 2020). In brief, the sequencing adaptors were removed from the pair-end reads, the bridge linker sequences were identified and the tags flanking the linkers were extracted. Tags identified (≥ 16 bp) were mapped to mm10 using BWA alignment and mem(Li and Durbin 2009) according to their tag length. The uniquely mapped, non-redundant pair-end tags (PETs) were classified as either inter-chromosomal (left tags and right tags aligned to the different chromosomes), intra-chromosomal (left tags and right tags aligned to the same chromosomes with genomic span > 8 kb) and self-ligation (left tags and right tags aligned to the same chromosomes with genomic span ≤ 8 kb) PETs. Interacting PETs (iPETs), the uniquely mapped, non-redundant pair-end tags (PETs) from both the inter- (left tags and right tags from different chromosomes) and intra-chromosomal (left tags and right tags with genomic span > 8 kb) PETs, were extended by 500 bp which was the average length of the sheared chromatin fragments. Multiple iPETs overlapping at both ends were then clustered as iPET-2, 3, … (clusters with 2, 3, … iPETs) to represent their interaction strength. Peaks were called using MACS (version 2.1.0.20151222) with *q* < 1E-8. The called peaks were used to determine interactions that are supported by the CTCF binding (interaction anchors must overlap at least 1 bp on a peak span).

### L. Enrichment and Network Analysis of Differentially Expressed Genes

Differential gene expression analysis was performed in R statistical software with R Studio(2020b; Team 2020b). *Rorb^h2/h2^* and *Rorb^h4/h4^* differentially expressed genes were analyzed individually. Those genes with FDR < 0.05 and p-value < 0.05 were uploaded to Mouse Mine, which cross-referenced several databases to identify EMAPA Anatomy Enrichment, Gene Ontology Biological Process, Reactome Pathways, and Mammalian Phenotype Ontology terms associated with genes overrepresented in each mutant set. Holm-Bonferroni correction for multiple testing was used consistently. The same list was uploaded to NetworkAnalyst3.0, and Generic PPI functionality was used to query STRING Interactome for evidence of protein-protein interactions with “require experimental evidence” selected and a default confidence score of 900(Szklarczyk *et al*. 2015; Zhou *et al*. 2019). The largest predicted subnetwork of associations, which contains *Rorb*, was arranged using the concentric circle layout with *Rorb* as the focal point, and a WalkTrap algorithm was used to explore functional modules. Functional annotations in the Reactome database were queried for each module of the top five modules ranked by p-value using built-in NetworkAnalyst3.0 functionality. *Rorb^h4/h4^* differentially expressed gene lists were cross-referenced with aggregated gene-disease association lists from OMIM and DisGeNET using the queries “Bipolar Disorder,” “Epilepsy,” and “Autism”(Pinero *et al*. 2017; 2020a). Gene sets were created in GeneWeaver containing the genes differentially expressed in each mutant compared to its wild-type control as well as lists of genes associated with each condition(Baker *et al*. 2012). Binary scores were added to indicate list membership. *Rorb^h4/h4^* GeneWeaver list ID: GS399094. Epilepsy: GS399098, ASD: GS399099, Bipolar Disorder: GS399100. The Jaccard similarly and Boolean Algebra tools were used with default settings to compute the degree of overlap of the *Rorb^h2/h2^* and *Rorb^h4/h4^* gene sets and those of the three neurological conditions. Disease gene sets were integrated with heterogeneous data from multiple species. The data contained gene sets from five web resources (GWASCatalog(Welter *et al*. 2014), OMIM(Amberger *et al*. 2015), MGI(Eppig *et al*. 2015), HPO(Kohler *et al*. 2014), DisGeNET(Pinero *et al*. 2017)). The list of differentially expressed genes and their average fold change between the *Rorb^h2/h2^* and *Rorb^h4/h4^* dataset were uploaded to Ingenuity Pathway Analysis software. The core pathway analysis with default settings was performed. The most significant network was exported. Expression levels and differential expression of genes that overlapped between *Rorb^h2/h2^* and *Rorb^h4/h4^* were visualized using Seaborn and NumPy and Python 3.0(Harris *et al*. 2020; Team 2020a). Cytoscape 3.8’s STRING Enrichment App using default parameters allowed assembly of interaction networks composed entirely of genes with *Rorb^h2/h2^* and *Rorb^h4/h4^* lists split on the basis of fold change direction (positive v. negative) using an established protocol published by Cytoscape. Nodes were organized using a Perfuse Force-Directed Layout with edge bundling (using default settings). Only the largest subnetworks are depicted in figures. Node size is scaled by the magnitude of fold-change and colored to indicate disease associations.

## RESULTS AND DISCUSSION

### Five mouse strains with gait and retinal phenotypes caused by spontaneous *Rorb* mutations

Between 2003 and 2020, five independent strains of spontaneous mutant mice were identified in vivaria at The Jackson Laboratory on the basis of their overt gait phenotypes **(Figure 1 A) (Video S1)**. The mice were originally referred to as “high-stepper,” but are denoted here as *Rorb^h1/h1^, Rorb^h2/h2^, Rorb^h3/h3^, Rorb^h4/h4^*, and *Rorb^h5/h5^*. All *Rorb* mutants displayed a pronounced gait phenotype with *Rorb^h5/h5^* being most dramatic. The lack of genetic complementation between intercrossed strains indicated the mutations were in the same gene. In all cases the frequency with which affected mice were born indicated recessive, monogenic inheritance. *Rorb^h3/h3^* mice displayed particularly poor viability and were necessarily omitted from subsequent experiments. *Rorb^h5/h5^* arose much later than the other mutants and was omitted from RNA sequencing experiments. Fine genetic mapping was performed by crossing affected (presumed homozygous) *Rorb^h1/h1^* mice on a C57BL/6J background with BALB/cByJ wildtype mice (see methods). This indicated that a 0.7Mb region of Chromosome 19, between markers D19MIT41 and D19MIT113, was associated with the gait phenotype. There are four protein coding genes within this interval: *Trpm6, Ostf1, Nmrk1*, and *RorB*(Eppig *et al*. 2015). The similarity between the gait phenotype of our mutants and that of previously described *Rorb* knockout mice suggested that *Rorb* mutations were causing the phenotype(Andre *et al*. 1998a).

**Figure 1.**
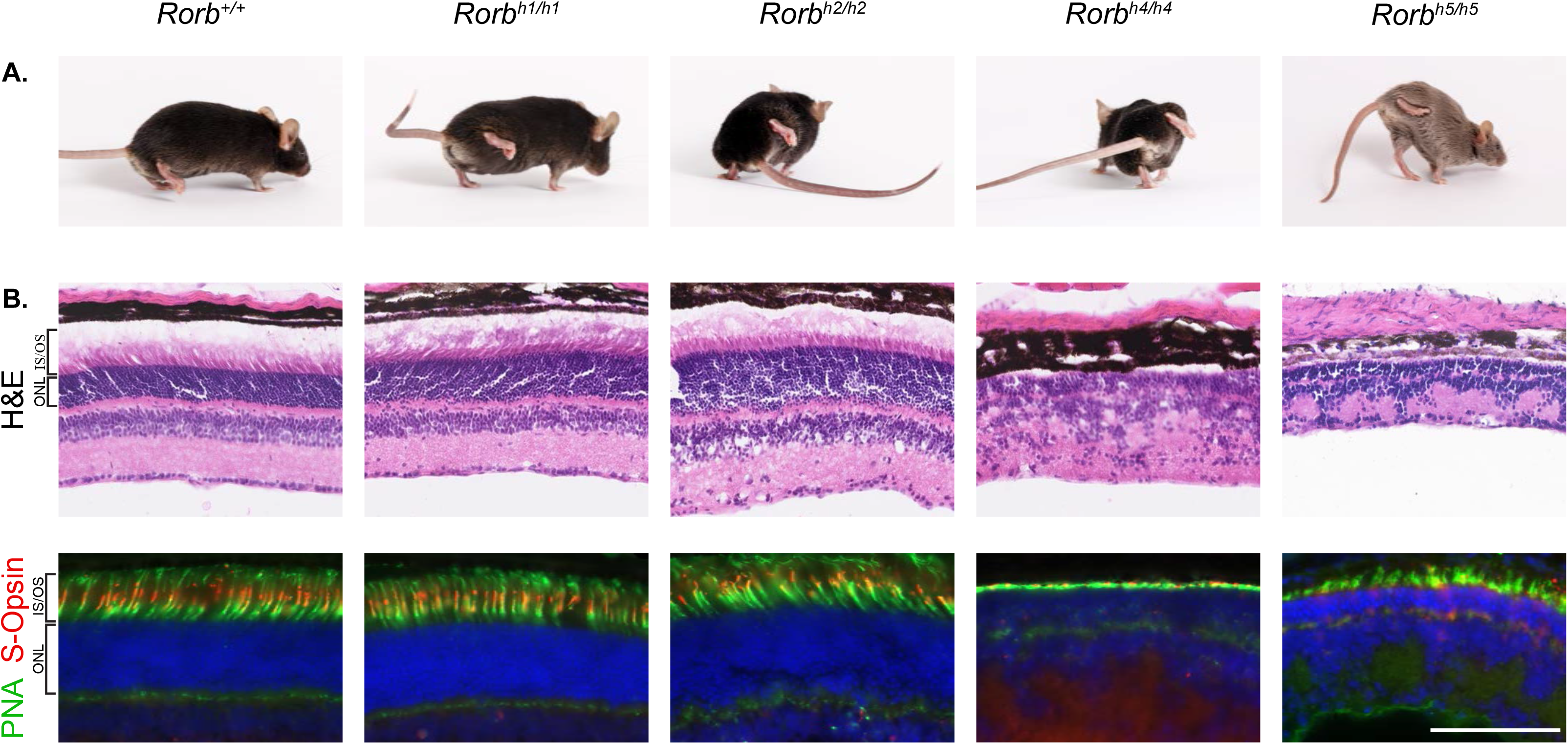
Gait Phenotypes and Retinal Histology in WT and *Rorb* mutant mice. *Rorb* mutant mice exhibit hindlimb hyperflexion with a distinctive arc during locomotion whereas WT mice never show this behavior. The height of abnormal limb elevation appears greatest in the *Rorb^h5/h5^* mutant with less deviation in *Rorb^h1/h1^, Rorb^h2/h2^*, and *Rorb^h4/h4^* (A). Top: Hematoxylin and Eosin-stained retinal sections imaged at 40X reveal structural changes in *Rorb^h4/h4^* and *Rorb^h5/h5^* mice where photoreceptor inner and outer segments are missing, and the retinal layers are highly disorganized. Bottom: Staining of retinal sections with peanut agglutinin (PNA) and S-Opsin demonstrates pronounced loss of inner and outer segments in *Rorb^h4/h4^* and *Rorb^h5/h5^* mice with other *Rorb* mutant mice showing no apparent changes in cone photoreceptors. Scalebar = 100mm, n=3 per genotype, 6 weeks of age (B).

In mice, *Rorb* consists of 11 exons spanning 181kb on chromosome 19. There are two isoforms distinguished by alternative first exons with independent initiation codons, which then splice to a common second exon **(Figure 2 A)**(Yates *et al*. 2020). PCR and Sanger sequencing of the two resultant transcripts, *Rorb1* and *Rorb2*, from *Rorb^h1/h1^* genomic DNA and cDNA revealed no deviation from the C56BL/6J reference sequences. For this reason, we included *Rorb^h1/h1^* in a large-scale project that sought to identify spontaneous mutations by whole-genome sequencing(Fairfield *et al*. 2015). This sequencing revealed a 326kb duplication impacting the 5’ end of *Rorb* that includes only the first exon of the *Rorb1* isoform **(Figure 2 B)**. *Rorb2* appears to be unaffected by the duplication. We were unable to identify the specific mutational events affecting *Rorb* in *Rorb^h2/h2^* and *Rorb^h3/h3^* mice, but the involvement of *Rorb* is supported by our complementation cross. Whole genome sequencing indicated an abnormally high density of reads aligned upstream of *Rorb* in both *Rorb^h1/h1^* and *Rorb^h2/h2^* mice, albeit in different patterns. This similarity suggests that the *Rorb^h2^* mutational event might be a duplication event similar to that of *Rorb^h1^* **(Figure S1 A).** Single nucleotide polymorphisms were identified in both *Rorb^h4/h4^* and *Rorb^h5/h5^* by Sanger sequencing. A single base pair change in *Rorb^h4/h4^* mice creates a premature stop codon in exon 8 of both *Rorb1* and *Rorb2* isoforms **(Figure 2 C)**. *Rorb^h5/h5^* mutants carry a thymine to cytosine transition in the ATG start codon of exon 1 of *Rorb1* **(Figure 2 D)**. This likely produces a *Rorb1* hypomorph as the next in-frame ATG occurs in exon 3 of both *Rorb* isoforms but should not affect *Rorb2*.

**Figure 2.**
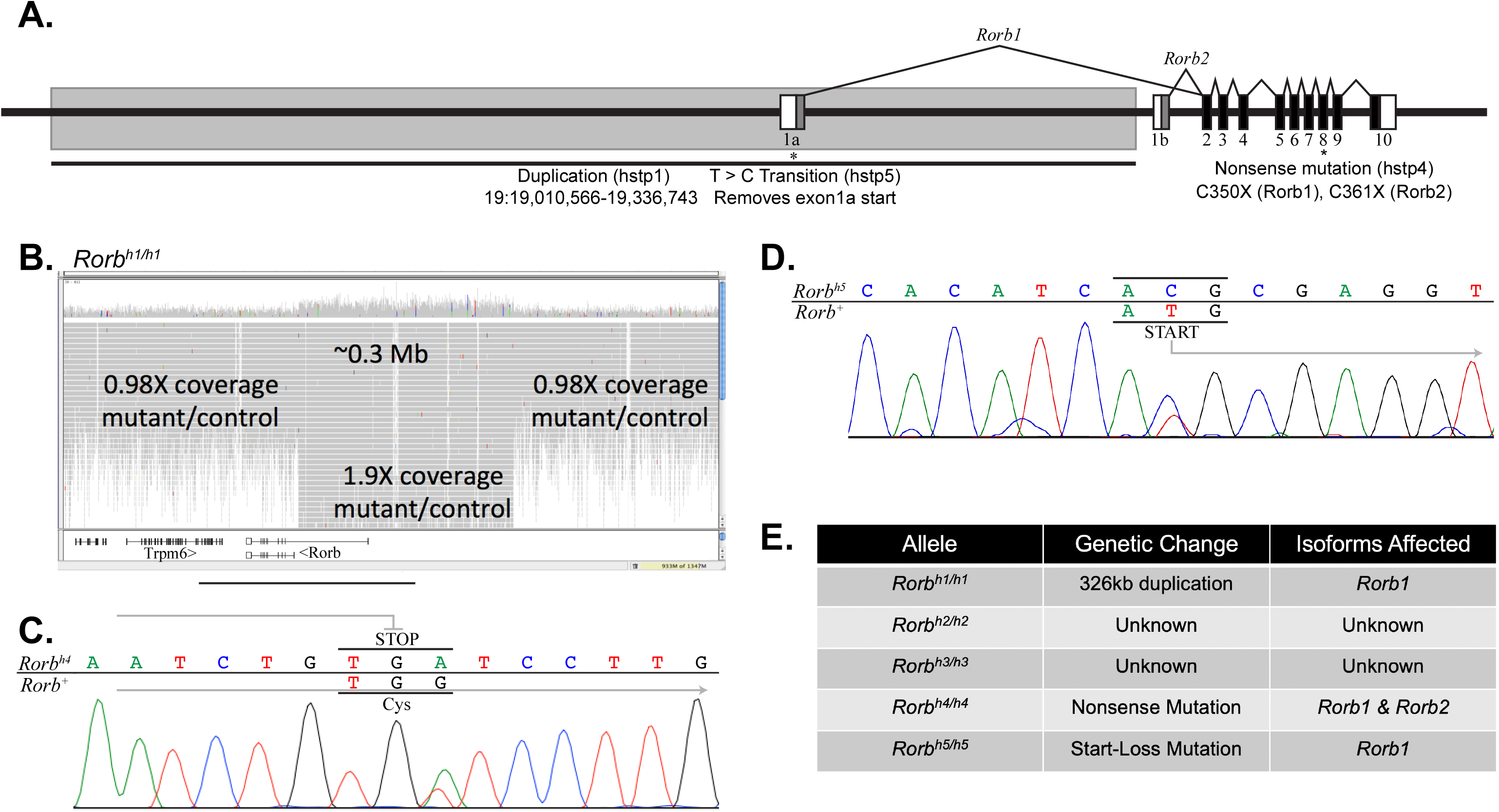
Identification of specific structural and sequence variation in *Rorb*. The *Rorb* locus encodes two isoforms, *Rorb1* and *Rorb2*. Each has 11 exons, and the two isoforms differ only in their usage of alternative 5’ first exons (exons 1a and 1b, respectively, shaded gray). Constitutive exons are represented by black bars while 5’ and 3’ untranslated regions are indicated by white bars. *Rorb* is located on the reverse strand of chromosome 19. The *Rorb^h1^* mutation is caused by a 326kb duplication (gray bar) encompassing the first exon of the *Rorb1* transcript. The *Rorb^h4^* mutation is a thymine to adenine transversion causing a nonsense mutation in exon 8 affecting both *Rorb1* and *Rorb2*. The *Rorb^h5^* mutation is a thymine to cytosine nonsynonymous substitution in the initiator methionine codon of *Rorb1* (A). The read depth of sequences mapped to the chromosome 19 region containing *Rorb* indicate 2X coverage compared to WT animals over a 326kb interval corresponding to the duplicated region in *Rorb^h1/h1^* mice. Note, the gene model is right to left because *Rorb* is encoded on the negative strand (B). The Sanger sequencing amplicon from a heterozygous *Rorb^+/h4^* animal reveals the T > A substitution in this line (C). The Sanger sequencing amplicon from a homozygous *Rorb^h5/h5^* mouse indicating the T > C substitution in this line (D). Table summarizing the mutation events in *Rorb* mutant mice (E).

Several previous articles describing *Rorb* null mice had demonstrated pronounced retinal phenotypes(Srinivas *et al*. 2006; Jia *et al*. 2009; Liu *et al*. 2013a; Fu *et al*. 2014). To investigate whether the retinas of our mutants display similar phenotypes, we performed staining with hematoxylin/eosin for gross morphology and immunofluorescence using peanut agglutinin (PNA) and anti-S-Opsin antibody to label all cones and only S-Opsin positive cones, respectively **(Figure 1 B)**. We only observed a strong retinal phenotype, one consistent with previous reports, in *Rorb^h4/h4^* and *Rorb^h5/h5^*. These mice exhibit clear thinning of the photoreceptor layer and prominent disorganization throughout the retinal nuclear layers. *Rorb^h1/h1^* and *Rorb^h2/h2^* have retinas that appear consistent with those of wildtype mice. Our mapping, the gait phenotype, and the observed retinal abnormalities support *Rorb* as the causal gene, and we next aimed to determine the genetic basis of phenotypic differences in these mutants.

### Changes in *Rorb* abundance caused by mutation

We next asked how expression from the *Rorb* locus was altered in mutants. We designed three primer sets, one specifically aligning to the *Rorb1* transcript, one specific to *Rorb2*, and one common to both isoforms. cDNA was derived from either whole brain **(Figure 3 A)** or retina **(Figure 3 B).** No data are provided for *Rorb2* expression in brain as we found no appreciable expression, consistent with previous findings(Schaeren-Wiemers *et al*. 1997; Andre *et al*. 1998b). Samples from seven genotypes were collected: *Rorb^h4/h4^, Rorb^+/h4^, Rorb^+/+^* (*Rorb^h4^* littermates), *Rorb^h1/h1^, Rorb^Cre/h1^, Rorb^Cre/Cre^, Rorb^+/+^* (*Rorb^Cre^* littermates). The *Rorb^Cre^* line(Harris *et al*. 2014) was generated by insertion of an IRES-Cre downstream of the translational stop with the intention of not disrupting *Rorb* expression. This line was included because homozygous *Rorb^Cre/Cre^* mice were reported to display “an abnormal walking gait with an exaggerated high step”, a description similar to the gait observed in our *high stepper* mice. In contrast to previous reports, however, we did not observe a gait phenotype in *Rorb^Cre/Cre^* mice, although compound heterozygous *Rorb^Cre/h1^* mice did display a gait phenotype, indicating that the *Rorb* locus is compromised by the insertion of IRES-Cre. Two different *Rorb^+/+^* cohorts were included to better control for potential background or litter effects, and transcript abundance for all groups was normalized to the average of these controls. *Rorb^h5/h5^* were not included in these experiments as they arose later.

**Figure 3.**
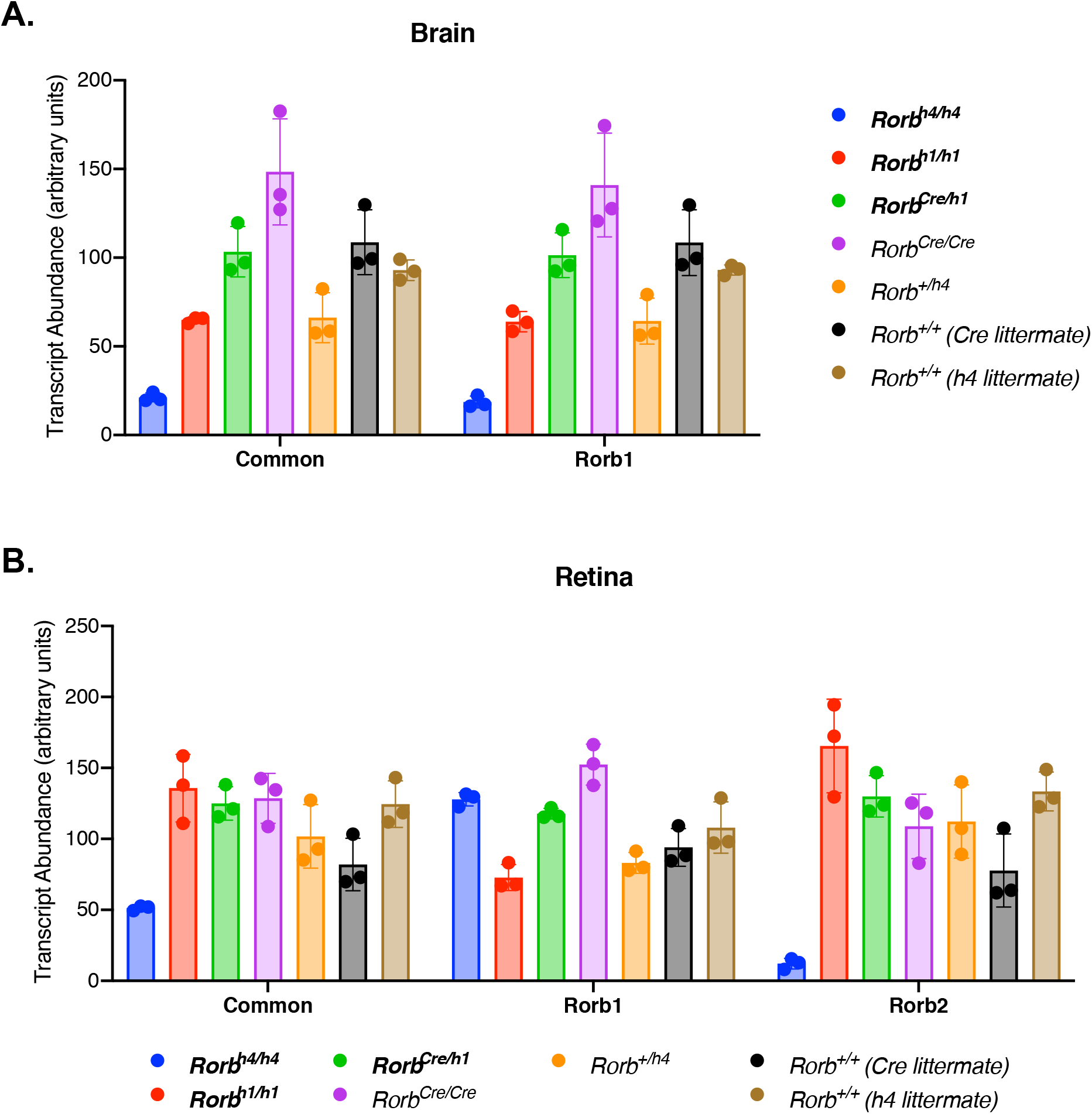
*Rorb* transcript abundance in retina and brain by qPCR. qPCR was performed on total RNA from brain (A) and retina (B) samples from animals of seven genotypes: *Rorb^h4/h4^, Rorb^+/h4^, Rorb^+/+^* (*Rorb^h4^* littermates), *Rorb^h1/h1^, Rorb^Cre/h1^, Rorb^Cre/Cre^*, and *Rorb^+/+^* (*Rorb^Cre^* littermates) (n = 3 per genotype). Primer sets that recognize both *Rorb1* and *Rorb2*, exclusively *Rorb1* and exclusively *Rorb2* were created. Results are normalized to expression of *Gapdh* and to the average of the two littermate control groups. Abundance is given as arbitrary units (equal to 100 times the fold change) and shown as mean ± standard deviation. Genotypes associated with the gait phenotype are listed in bold. Results are presented from mice aged 20-25 days. *Rorb^h4^* heterozygous and homozygous mice were compared to their WT littermates. *Rorb^Cre^*-carrying mice were compared to their WT littermates. *Rorb^h1/h1^* mice were compared to both WT groups. Black asterisks indicate significance compared to WT *Cre* littermates, and brown asterisks indicate significance compared to WT *h4* littermates. Data were analyzed by ordinary one-way ANOVA with Tukey’s multiple comparisons. ****p<0.0001, ***p<0.001, **p<0.01, *p<0.05.

We hypothesized a general correlation between phenotypic severity and the abundance of *Rorb* transcript. In line with our expectations, *Rorb^h4/h4^* cDNA contained the lowest amount of *Rorb* transcript in both brain and retina, presumably due to nonsense-mediated decay (NMD). Interestingly, unlike *Rorb2, Rorb1* was present at WT levels in the *Rorb^h4/h4^* retina, suggesting that perhaps the transcript is not effectively degraded by NMD. *Rorb^h1/h1^* samples show ~40% decrease in *Rorb1* abundance relative to WT, while *Rorb2* is unaffected. This is consistent with the identified duplication in these mice, which impacts only the *Rorb1* isoform, and the fact that *Rorb2* is expressed in retina, but not in brain. Functional redundancy of the two isoforms in the retina, combined with an observed increase in *Rorb2 levels*, may explain why these animals do not display the retinal phenotype. This experiment also yielded additional surprising results. First, compound heterozygous *Rorb^Cre/h1^* mice, which exhibit the abnormal gait, display *Rorb* transcript abundance similar to that of WT mice. Second, *Rorb^+/h4^* mice do not exhibit the gait but have levels of *Rorb* similar to those of *Rorb^h1/h1^* mice. Cumulatively, these findings suggest that while levels of *Rorb* transcript roughly correlate with phenotypic severity, they are not predictive.

### *Rorb* expression patterns in brain and spinal cord

We next examined the expression pattern of *Rorb*. Based on the long-standing use of *Rorb* as a marker of layer IV neurons in the somatosensory cortex, we examined this population of cells using the *Rorb^Cre^* driver line crossed to the *ROSA^tdT^* reporter line. In order to view *Rorb*-expressing cells in the context of the high stepping phenotype, mice carrying the driver and reporter were crossed to *Rorb^h1/h1^* animals, resulting in compound heterozygous *Rorb^Cre/h1^* mice, which exhibit the gait phenotype. In confocal images of coronal sections of primary somatosensory cortex, both *Rorb^+/Cre^; ROSA^tdT^* and *Rorb^Cre/h1^; ROSA^tdT^* samples had a brightly-labeled band of cells in the predicted cortical layer IV and stretching across the full domain occupied by the somatosensory cortex **(Figure 4 A & B)**. The number and distribution of cells was indistinguishable between the two genotypes.

**Figure 4.**
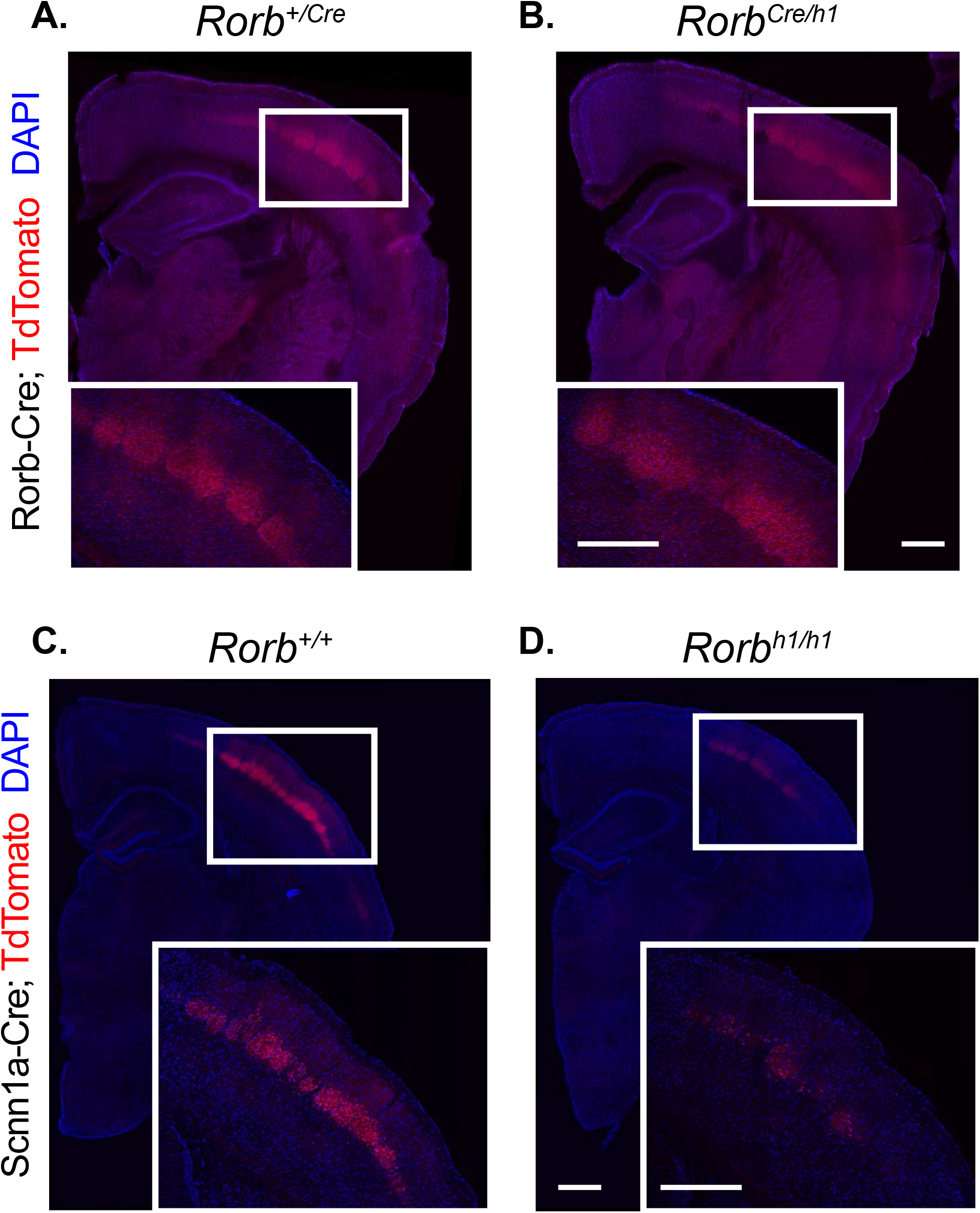
Changes in reporter expression in layer 4 somatosensory cortex of *Rorb^hstp^* mice. *Rorb^+/Cre^; ROSA^tdT^* and *Rorb^Cre/h1^; ROSA^tdT^* cortical sections imaged for tdTomato reveal similar numbers and intensity of positive cells within layer 4 of somatosensory cortex (A,B). Levels of the tdTomato reporter driven by *Scnn1a^Cre^* did differ by genotype, however (C,D). Markedly fewer positive cells were seen in *Scnn1a^+/Cre^; Rorb^h1/h1^; ROSA^tdT^* tissue compared to control. Scale bar = 500 um.

To further investigate the fate of *Rorb*-expressing cells in *Rorb* mutants, we used the *Scnn1a-Cre* driver line(Harris *et al*. 2014) in combination with the *ROSA^tdT^* reporter and the *Rorb^h1^* mutation. Both *Scnn1a^+/Cre^; Rorb^+/+^; ROSA^tdT^* and *Scnn1a^+/Cre^; Rorb^h1/h1^; ROSA^tdT^* samples labeled a population of spiny excitatory neurons in layer IV of the somatosensory cortex **(Figure 4 C & D)**; however, the signal was significantly brighter in WT versus mutant tissue due to a smaller number of labeled cells in the mutant animal. This decrease in labeled cells could either reflect changes in terminal differentiation or more specific effects on the expression of the reporter.

Given the expression of *Rorb* in sensory areas, the gait phenotype, and the above findings of changes in somatosensory cortex, we further examined sensory nerves and sensory-related behavior in *Rorb^h1/h1^* mice. We found no differences in axon number or area in femoral nerve motor and sensory branches of mutants compared to heterozygous littermates, which do not exhibit the gait phenotype **(Figure S2 A-B)**. A modest significant decrease in average area of the entire nerve was seen for the femoral motor, but not the sensory, nerve **(Figure S2 C)**. No significant differences between genotypes were detected on either the hot plate test of thermal nociception or the von Frey test of mechanosensation **(Figure S2 D & E)**. Thus, we do not detect global changes in peripheral nerve anatomy or sensory capability.

Given the prominent gait phenotype of the mutant mice and previous explanation of the involvement of *Rorb* expression in inhibitory interneurons of the spinal cord and the gait phenotype(Koch *et al*. 2017), we next examined this tissue using the *Rorb^Cre^; ROSA^tdT^* reporter system described above in both high-stepping *Rorb^Cre/h1^* animals and unaffected controls. Confocal imaging of the tdTomato reporter revealed expression of *Rorb* in multiple regions of the spinal cord, including the dorsal horn and scattered cells within the white matter **(Figure S3 A & B)**. No differences were observed between *Rorb^+/Cre^; ROSA^tdT^* and *Rorb^Cre/h1^; ROSA^tdT^* samples. To determine which layers of the dorsal horn contain *Rorb*-positive fibers, we performed co-labeling with CGRP and isolectin-b4 **(Figure S3 C & D)**, which label peptidergic and nonpeptidergic nociceptive fibers, respectively(Zeilhofer *et al*. 2012). We saw overlap of the tdTomato reporter with isolectin b4, but not CGRP, suggesting involvement of *Rorb*-expressing cells in the nonpeptidergic nociception pathway. Again, no differences were observed between *Rorb* mutant mice and controls. We also performed immunohistochemistry for myelin basic protein and found colocalization with the tdTomato reporter in white matter in both WT and *Rorb* mutants, suggesting that some *Rorb*-positive cells in the spinal cord are oligodendrocytes **(Figure S3 E & F).**

### RNA sequencing reveals transcriptomic changes and interaction of disease-associated gene products

To better understand the effects of *Rorb* mutations on gene expression, we performed RNAseq on samples of whole brain from WT, *Rorb^h1/h1^, Rorb^h2/h2^*, and *Rorb^h4/h4^* mice (n = 3 per genotype at 6 weeks of age). When read depth mapping to the genomic territory surrounding *Rorb* is compared among groups, dramatic differences are apparent **(Figure S1 A)**. To examine the impact of the *Rorb^h1^* duplication on chromatin interactions around *Rorb*, we used chromatin immunoprecipitation and paired end sequencing (ChIA-PET) of neural progenitor cells from the developing brain at E12.5-14.5. This approach allowed us to map CTCF-mediated chromatin interactions across the duplicated interval. We found that compared to WT, chromatin interactions were either completely missing or were rearranged in Rorb^h1/h1^ mutant NPCs. Missing interactions included interactions between the *Rorb* locus and the neighboring gene *Trpm6*, as well as a strong loop in the intergenic region upstream of Rorb. By comparing these chromatin interaction data with publicly available chromatin state maps of mouse development, we found that this missing loop overlaps with a strong distal enhancer that is specific to the developing forebrain at E14.5-16.5. Such enhancers have been identified through the intersection of ENCODE3 chromatin marks (ChromHMM) with known topologically associated domains (TAD’s) and EPDnew mouse promoters from the Eukaryotic Promoter Database(Gorkin *et al*. 2020). While more work is needed to experimentally validate this enhancer and to characterize the impact of this disrupted enhancer interaction on developmental expression of *Rorb*, it suggests that the *Rorb^h1^* duplication is likely to disrupt the spatiotemporal developmental regulation of this gene during development (**Figure S1 B**). While there is little transcription ~400kb upstream of *Rorb* in WT and *Rorb^h4/h4^* mice, a consistent expression pattern is noted in both *Rorb^h1/h1^* and *Rorb^h2/h2^* animals, though these patterns differ from one another. *Rorb^h1/h1^* samples have a broad distribution of reads throughout much of the duplicated interval, whereas *Rorb^h2/h2^* samples show fewer reads overall, mostly concentrated ~350kb upstream of the first exon of *Rorb1*. These results likely indicate that, in *Rorb^h1/h1^* mice, transcription begins normally at the duplicated upstream *Rorb1* promoter, but then abnormal transcripts are produced that include sequence from the duplicated region. Despite this altered pattern of expression, 5’-RACE indicates that normal *Rorb1* transcript is produced in *Rorb^h1/h1^* mice, and no other major species were detected using this approach **(Figure S1 C)**. The presence of aberrant reads in *Rorb^h2/h2^* mice suggests that a similar structural change may have occurred in the genome of these animals, though to date we have not been able to identify such an event.

To better understand of the transcriptome-wide effects of the mutant transcription factor *Rorb*, we performed differential gene expression analysis in *Rorb^h1/h1^, Rorb^h2/h2^*, and *Rorb^h4/h4^* mice using a p-value <0.05 and a false-discovery rate (FDR) <0.05 cutoff **(Spreadsheet S1)**. The only gene whose expression was significantly changed in all three mutants compared to WT was *Rorb* itself. In *Rorb^h1/h1^* mutants, the only differentially expressed gene aside from *Rorb* was *SoxOs1*, a long non-coding RNA gene. Both *Rorb* and *SoxOs1* had negative fold-changes in mutants. *Rorb^h2/h2^* had a total of 149 genes with altered expression (98 positive and 51 negative log2 fold changes (log2FC) relative to WT), and *Rorb^h4/h4^* had 524 genes (318 positive and 206 negative log2FC genes relative to WT) with 33 differentially expressed genes overlapping between *Rorb^h2/h2^* and *Rorbh^4/h4^* (**Figure 5 A**).

**Figure 5:**
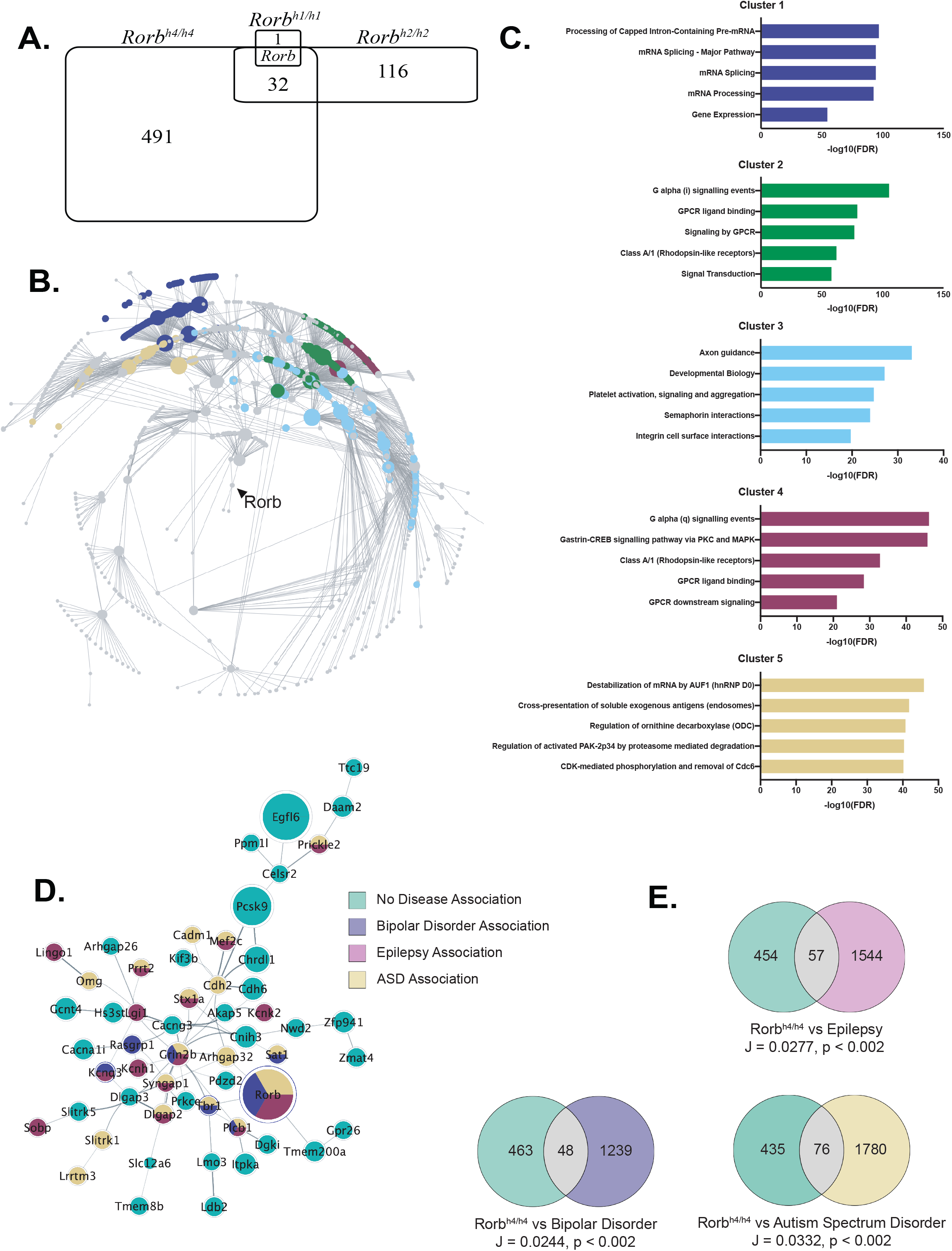
Differential expression and gene set enrichment analysis of *Rorb^h4/h4^* mice.. RNAseq data were also analyzed globally to determine the number of differentially expressed genes in each mutant compared to control. Including *Rorb*, the 33 differentially expressed genes in common between *Rorb^h2/h2^* and *Rorb^h4/h4^* (A). An unordered concentric-circle STRING Interactome network of molecular interactions with RORB as focal protein and WalkTrap clustering reveals those gene modules that are likely impacted by *Rorb* mutation. The five most significant WalkTrap functional modules with are color coded (B) and match the functional ontology terms associated with each module (log10 p-values) (C). Cytoscape STRING protein-protein interaction networks of genes with negative logFC values reveal high centrality of numerous disease-associated genes. Node size is scaled by relative logFC and colored to indicate disease involvement (D). Jaccard comparison of *Rorb^h4/h4^* differentially expressed gene lists with disease-associated gene sets from OMIM and DisGeNet reveals significant overlap (E). Animals were 6 weeks old, n=3 per genotype.

We used Mouse Mine to search enrichment of Reactome Pathways, EMAPA anatomy terms, Gene Ontology Biological Processes (GO:BP), Gene Ontology Molecular Functions (GO:MF), and Mammalian Phenotype Ontology (MPO) terms with lists of differentially expressed genes(Bult *et al*. 2019). Enrichment of MPO terms related to nervous system phenotypes and morphology were observed for differentially expressed genes in *Rorb^h2/h2^* (**Table S1**). Analysis of enriched MPO terms for *Rorb^h4/h4f^* revealed a variety of ontologies related to abnormal behavioral phenotypes, glutamatergic signaling, locomotion, and body size (**Table S2**). When the list of differentially expressed genes from *Rorb^h4/h4^* mutants was separated by the direction of log2FC (positive v. negative) it became apparent that bodyweight-related annotations were identified due to genes with positive log2FC while those related to the nervous system were due mostly to genes with negative log2FC (**Table S3**). Importantly, those genes with negative log2FC are significantly associated with the Reactome Pathway leucine-rich glioma inactivated / a disintegrin and metalloproteinase family (LGI-ADAM) interactions (Holm-Bonferroni p = 0.0185). LGIs are secreted synaptic proteins containing a leucine-rich repeat and epilepsy-associated domains that interact with cell surface receptors (ADAM proteins) and Syntaxin1 during synapse development and myelination(Gu *et al*. 2002; Staub *et al*. 2002; Kegel *et al*. 2013).

We next assembled a protein-protein interaction network based on differentially expressed genes from *Rorb^h4/h4^* with the STRING Interactome database using parameters set to require experimental evidence and a confidence value of 900 or greater with NetworkAnalyst3.0(Zhou *et al*. 2019). We visualized the largest predicted subnetwork using a concentric circle layout taking *Rorb* as the focal point and identified modules within the network using a WalkTrap algorithm **(Figure 5 B)**. Forty-seven modules in total were identified with p-value < 0.05, and we chose to investigate the top five modules ranked by p-value. Reactome terms associated with the genes underlying each of the top modules are varied **(Figure 5 C)**, and contain numerous terms involving cellular adhesion, the actin cytoskeleton, axon guidance, and protein folding. The term “rhodopsin-like receptor” appears in two of the top five modules. Though not one of the top five modules, Rorb and several other steroid-thyroid hormone-retinoid receptors are part of a module (p = 1.45e-16) associated with Reactome terms related to transcription and gene expression, which may mediate effects downstream of *Rorb* disruption.

We then cross-referenced genes differentially expressed in *Rorb^h4/h4^* with lists of disease-associated genes from the DisGeNet and OMIM databases(Pinero *et al*. 2017; 2020a) for the neurodevelopmental conditions with which *Rorb* has previously been associated, namely autism spectrum disorder (ASD), epilepsy, and bipolar disorder(Mansour *et al*. 2009a; McGrath *et al*. 2009; McCarthy *et al*. 2012; Lai *et al*. 2015; Rudolf *et al*. 2016; Sadleir *et al*. 2020; Satterstrom *et al*. 2020). STRING protein-protein interaction (PPI) networks, split on the basis of the direction of differential gene expression in *Rorb^h4/h4^* mice relative to WT (positive v. negative), were created in Cytoscape 3.8, and disease-association of individual nodes are indicated **(Figure 5 D)**. Disease associated nodes exhibit relatively high betweenness centrality in the network composed of those genes with decreased expression in *Rorb^h4/h4^*. Approximately 5% of ASD-related genes identified in a large human ASD GWAS(Satterstrom *et al*. 2020) have mouse orthologues amongst the 25 PPI components most closely associated with *Rorb* by number of shared edges in our differential gene expression data(Satterstrom *et al*. 2020). For each condition, approximately 10% of the differentially expressed genes from *Rorb^h4/h4^* had an association with each neurological condition **(Figure 5 E)**, and Jaccard Analysis performed using GeneWeaver indicated these overlaps were significant(Baker *et al*. 2012).

It must be noted that the *Rorb^h4/h4^* mutation arose on a C57BL/6J congenic carrying a mutation in the *Il1r1* gene. We performed an analysis of Gene Ontology annotation overrepresentation for the entire set of differentially expressed genes in this mutant and did not see an enrichment of immune-related terms, or other pathways associated with *Il1r1* mutation such as “IL-1 signaling pathway” or “Bacterial infections in CF airways”(Kanehisa and Goto 2000; Rouillard *et al*. 2016; Stelzer *et al*. 2016). This leads us to believe that the majority of ontology terms detected in our analysis result from *Rorb* mutation. We also found no hub protein interactions and only one protein-interaction partners of *Il1r1* from the Pathway Commons database amongst differentially expressed genes (1 of 524 genes) in *Rorb^h4/h4^* (Rouillard *et al*. 2016), and no GAD Gene-Disease, GEO Disease Signatures, GWASdb SNP associations between *IL1R1* and epilepsy, autism spectrum disorder, or bipolar disorder(Becker *et al*. 2004; Rouillard *et al*. 2016). It is worth noting that *Il1r1* knockout has been associated with reduced susceptibility to develop seizures after induced febrile seizures in young mice(Feng *et al*. 2016). The authors demonstrate this resistance occurs through upregulation of endocannabinoid signaling, which we do not observe in our dataset. Therefore, we suspect transcriptional changes in *Rorb^h4/h4^* brain result from changes in *Rorb* expression and function, though we encourage further study of these transcriptional networks before speculating on therapeutic targets or drawing clinically-actionable insights.

We next examined the more selective set of 33 genes that are differentially expressed in both *Rorb^h2/h2^* and *Rorb^h4/h4^* mice relative to wildtype controls. Differentially expressed genes shared between mutants generally exhibit the same direction of differential expression relative to wildtype, and in most cases the magnitude of the log fold-change was greater in *Rorb^h4/h4^* compared to *Rorb^h2/h2^* (**Figure 6 A**). Mouse Mine was used to examine enrichment of process terms and pathways(Motenko *et al*. 2015). Both GO:BP and GO:MF terms indicate a change in protein folding activities and an overrepresentation of genes whose products are involved in pathways associated with heat shock and stress **(Figure 6 B).** Subjecting this set of genes to Ingenuity Pathway Analysis using default settings led to the identification of an interaction network that includes 13 of the genes from our set with a p-value of 10e-32 **(Figured 6 C)**. This set is enriched for the biological functions: cellular assembly and organization, cellular development, and cellular growth and proliferation. A literature review indicated that 36% (12/33) of these shared genes have been investigated on the basis of their association with epilepsy, 33% (11/33) for involvement in ASD, and 25% (9/33) for involvement in bipolar disorder (**TABLE 1**). These shared genes may represent drivers of transcriptional remodeling that takes place in *Rorb* mutants or markers of *Rorb*-related neurodevelopmental disruption.

**Figure 6:**
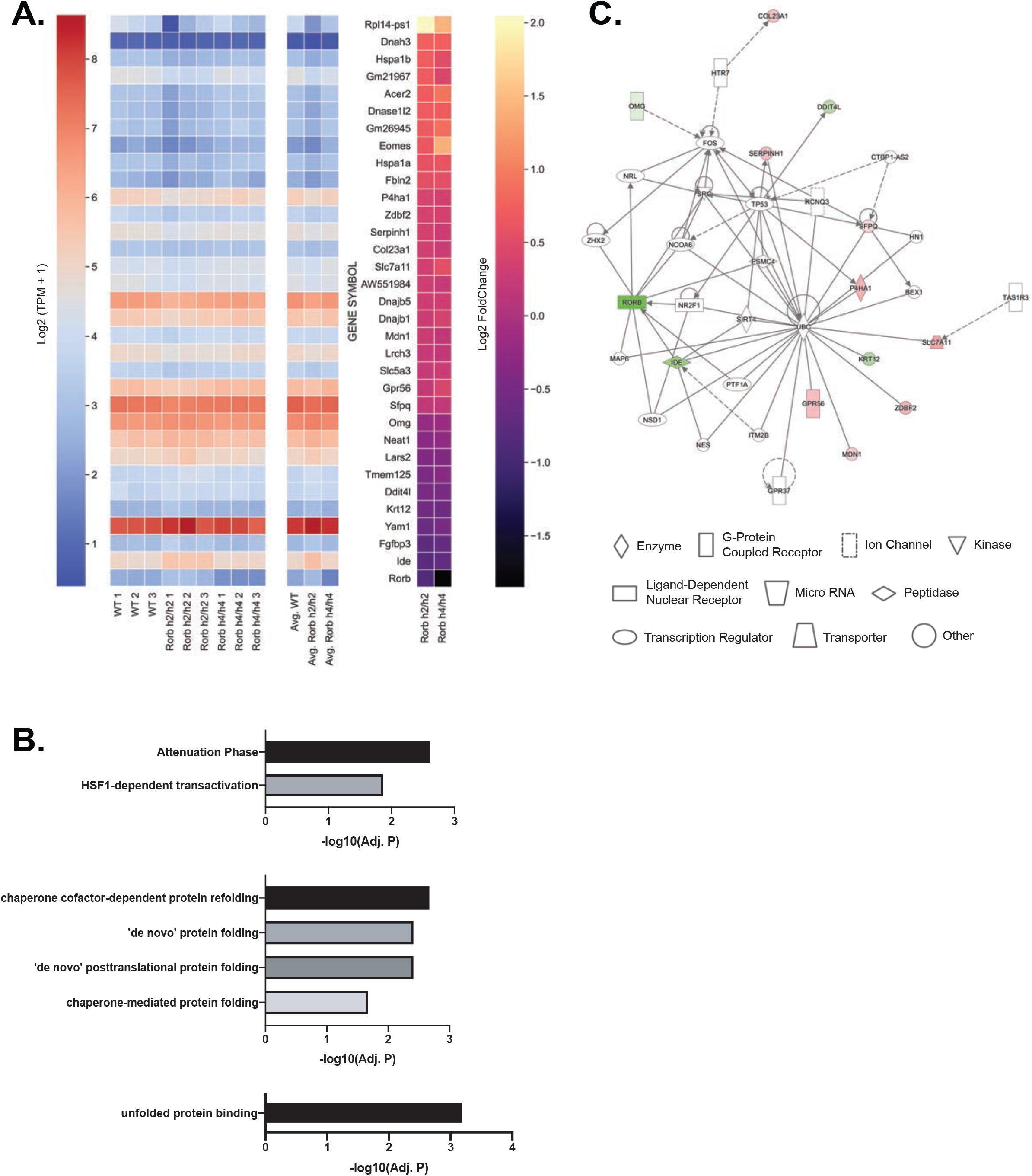
Biological processes, pathways, and networks enriched in *Rorb^h4^* differentially expressed gene list. A heatmap of log2 (TPM +1) by sample (left), averaged across samples (middle), and logFC of each group relative to wildtype (right) reveals comparable expression and consistent direction of differential expression in each mutant group (A). Ontology query of the overlapping gene set on Mouse Mine revealed pathway, gene ontology process and function terms related to protein folding and chaperone proteins (B). This overlapping gene set was examined by Ingenuity Pathway Analysis and the most significant pathway, containing 13 of the genes in the set, is shown. Green indicates downregulation of the gene, red indicates upregulation (C). Animals were 6 weeks old, n=3 per genotype.

**Table 1:**
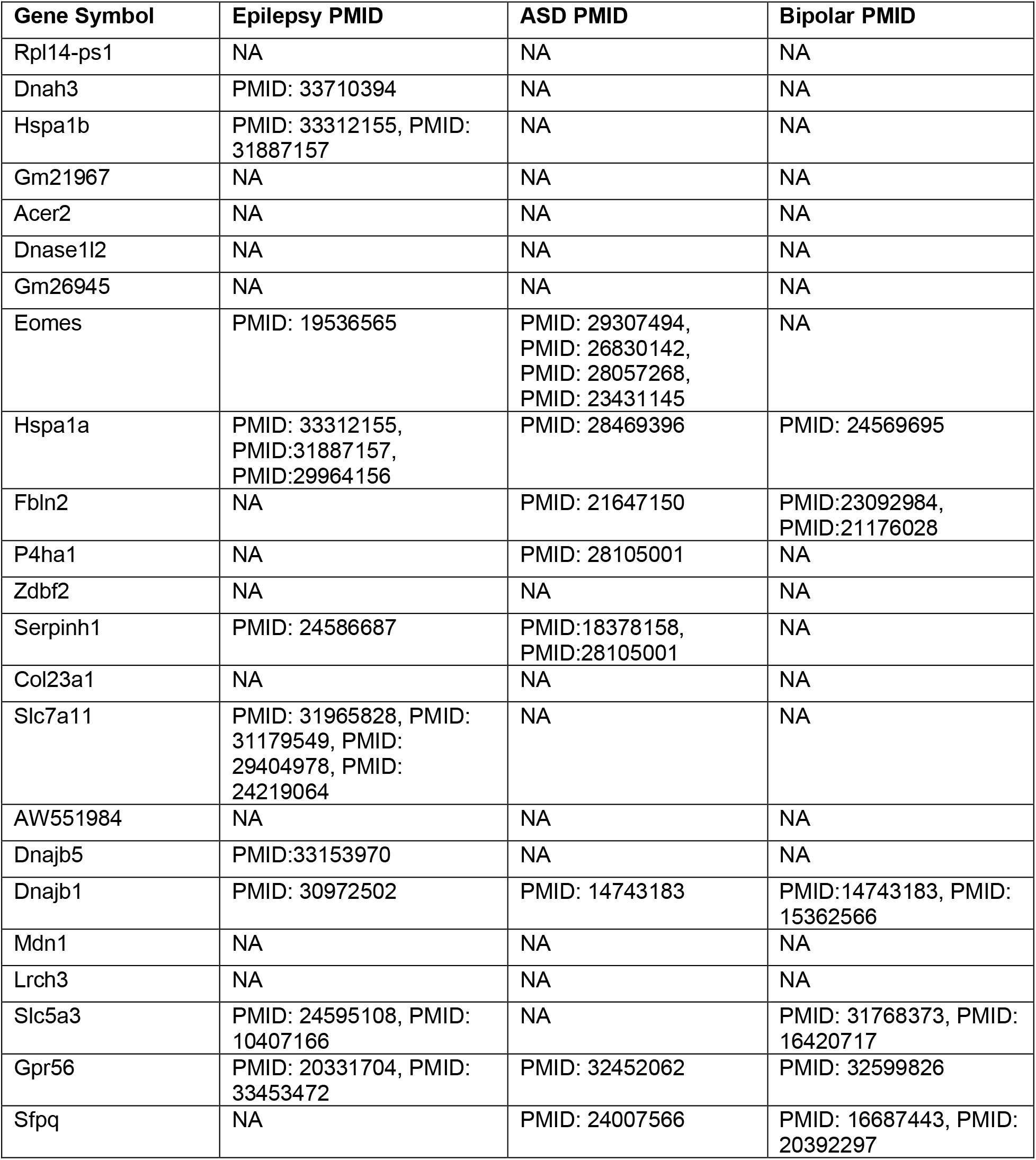
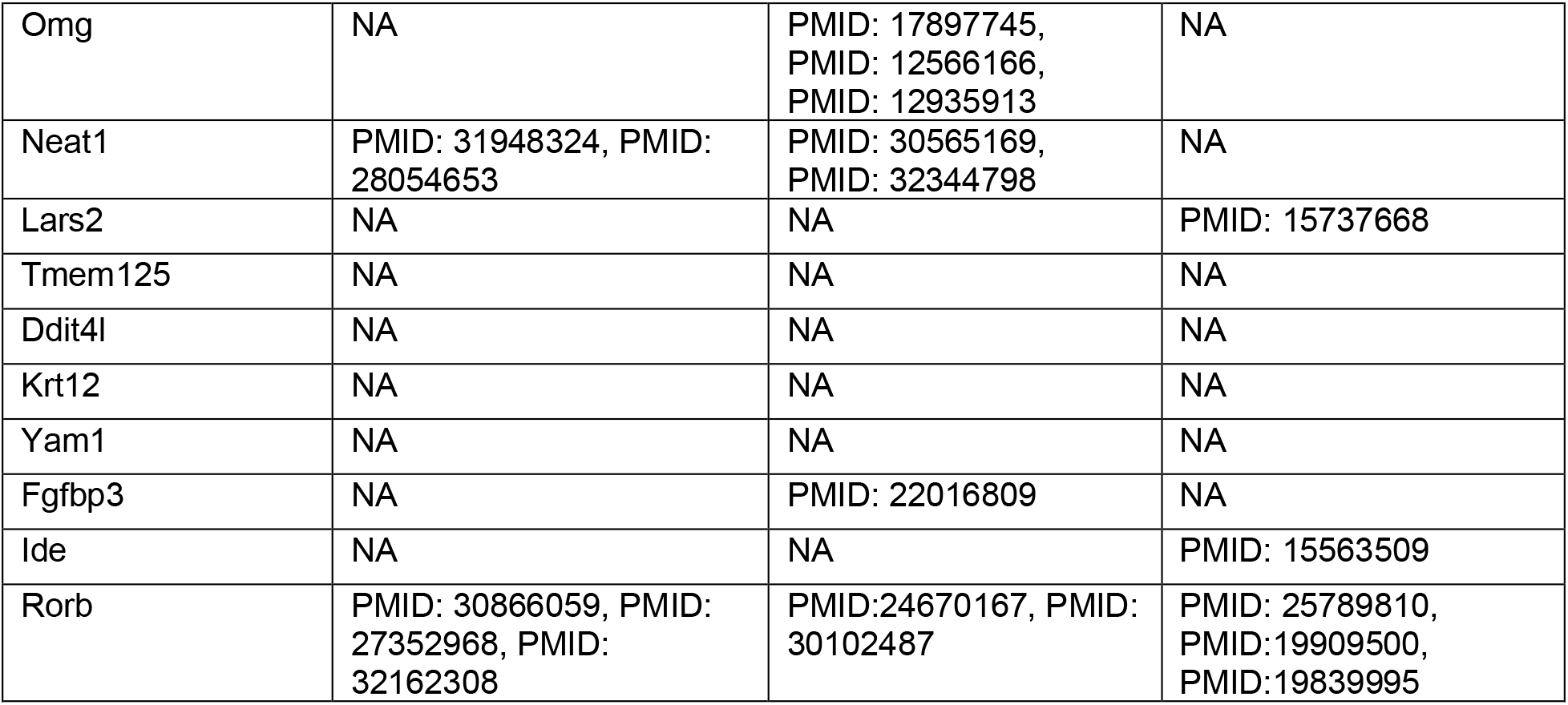
Rorb^h2/h2^ and Rorb^h4/h4^ Overlapping Differentially Expressed Genes Disease Associations.

| Gene Symbol | Epilepsy PMID | ASD PMID | Bipolar PMID |
| --- | --- | --- | --- |
| Rpl14-ps1 | NA | NA | NA |
| Dnah3 | PMID: 33710394 | NA | NA |
| Hspa1b | PMID: 33312155, PMID: 31887157 | NA | NA |
| Gm21967 | NA | NA | NA |
| Acer2 | NA | NA | NA |
| Dnase1l2 | NA | NA | NA |
| Gm26945 | NA | NA | NA |
| Eomes | PMID: 19536565 | PMID: 29307494, PMID: 26830142, PMID: 28057268, PMID: 23431145 | NA |
| Hspa1a | PMID: 33312155, PMID:31887157, PMID:29964156 | PMID: 28469396 | PMID: 24569695 |
| Fbln2 | NA | PMID: 21647150 | PMID:23092984, PMID:21176028 |
| P4ha1 | NA | PMID: 28105001 | NA |
| Zdbf2 | NA | NA | NA |
| Serpinh1 | PMID: 24586687 | PMID:18378158, PMID:28105001 | NA |
| Col23a1 | NA | NA | NA |
| Slc7a11 | PMID: 31965828, PMID: 31179549, PMID: 29404978, PMID: 24219064 | NA | NA |
| AW551984 | NA | NA | NA |
| Dnajb5 | PMID:33153970 | NA | NA |
| Dnajb1 | PMID: 30972502 | PMID: 14743183 | PMID:14743183, PMID: 15362566 |
| Mdn1 | NA | NA | NA |
| Lrch3 | NA | NA | NA |
| Slc5a3 | PMID: 24595108, PMID: 10407166 | NA | PMID: 31768373, PMID: 16420717 |
| Gpr56 | PMID: 20331704, PMID: 33453472 | PMID: 32452062 | PMID: 32599826 |
| Sfpq | NA | PMID: 24007566 | PMID: 16687443, PMID: 20392297 |
| Omg | NA | PMID: 17897745,<br>PMID: 12566166,<br>PMID: 12935913 | NA |
| Neat1 | PMID: 31948324, PMID:<br>28054653 | PMID: 30565169,<br>PMID: 32344798 | NA |
| Lars2 | NA | NA | PMID: 15737668 |
| Tmem125 | NA | NA | NA |
| Ddit4l | NA | NA | NA |
| Krt12 | NA | NA | NA |
| Yam1 | NA | NA | NA |
| Fgfbp3 | NA | PMID: 22016809 | NA |
| Ide | NA | NA | PMID: 15563509 |
| Rorb | PMID: 30866059, PMID:<br>27352968, PMID:<br>32162308 | PMID:24670167, PMID:<br>30102487 | PMID: 25789810,<br>PMID:19909500,<br>PMID:19839995 |

**Table 2:** Mouse strains used in this study.

| Official Strain Designation | Abbreviation |
| --- | --- |
| B6.Cg-Gt(ROSA)26Sor<sup>tm14(CAG-tdTomato)Hze/J</sup> | *ROSA<sup>tdT</sup>* |
| B6;129S-*Rorb*<sup>tm1.1(cre)Hze/J</sup> | *Rorb<sup>Cre</sup>* |
| B6;C3-Tg(Scnn1a-cre)3Aibs/J | *Scnn1a<sup>Cre</sup>* |
| C57BL/6J-*Rorb*<sup>hstp/J</sup> | *Rorb<sup>h1</sup>* |
| B6.Cg-Tyr<sup>c-2J</sup> *Rorb*<sup>hstp-4J</sup>/GrsrJ | *Rorb<sup>h2</sup>* |
| B6.Cg-Tyr<sup>c-2J</sup> *Rorb*<sup>hstp-3J</sup>/GrsrJ | *Rorb<sup>h3</sup>* |
| B6.Cg-*Rorb*<sup>hstp-2J</sup>/GrsrJ | *Rorb<sup>h4</sup>* |
| DBA/1J-*Rorb*<sup>hstp-5J</sup>/Rwb | *Rorb<sup>h5</sup>* |

## CONCLUSION

In this study, we present an allelic series of spontaneous mutations in *Rorb*. We were able to identify the causative mutation in three out of five of these mice: the *Rorb^h1^* allele is a duplication of 326kb impacting only the first exon of the *Rorb1* transcript, the *Rorb^h4^* allele is a nonsense mutation in exon 8, which is common to both transcripts, and the *Rorb^h5^* allele contains a thymine to cytosine transition in the ATG start codon of the first exon of the *Rorb1* transcript. In addition to the gait phenotype by which all mutants were identified, we also observed an absence of cone photoreceptors *in Rorb^h4/h4^* and *Rorb^h5/h5^* mutants. We also describe an expression pattern of *Rorb* in sensory areas, including the spinal cord, thalamus, and somatosensory cortex, consistent with previous studies(Koch *et al*. 2017; Clark *et al*. 2020). We have also demonstrated that, although the number of *Rorb*-expressing cells in somatosensory cortex is not dramatically different between mutant and WT mice, the expression of *Scnn1a* is markedly different, with many fewer cells labeled in mutant mice. We therefore support the finding that, in layer 4 neurons, *Rorb* is important for cell fate and/or terminal differentiation, similar to what has been reported previously regarding cortical barrels, spinal cord, and retina (Kautzmann *et al*. 2011; Liu *et al*. 2013b; Fu *et al*. 2014; Wang *et al*. 2014; Clark *et al*. 2020; Carneiro *et al*. 2021). Thus, while *Rorb*-expressing cells are still present in sensory areas in mutant mice, they may not function normally in cortical circuitry.

It is intriguing that a relatively subtle mutational event, the duplication of the first exon, in *Rorb^h1/h1^* mice can result in the obvious gait phenotype displayed by these animals. 5’-RACE experiments detected the presence of WT transcript, and qPCR suggests that the levels of *Rorb1* are only decreased by ~40% in the adult brain. This decrease is similar to that seen in *Rorb^+/h4^* heterozygous animals, which have a normal gait. These data, in combination with the lack of coding mutations in three out of our five mutants, suggest that either: 1) the expression of the *Rorb* gene is highly sensitive to any mutational changes, or 2) RORB protein function is highly sensitive to levels, developmental timing, or pattern of expression. The fact that the gait phenotype is apparent from the time the animals start walking and does not worsen with age would be consistent with *Rorb* misregulation during development, and our chromatin interaction data from neural progenitor cells shows that the duplication disrupts a putative developmental enhancer interaction. While no coding mutations were detected in *Rorb^h2/h2^* mice, our RNAseq data suggest that a structural change upstream of *Rorb*, similar to that in the *Rorb^h1^* allele, may be to blame. In addition to the surprising sensitivity of the *Rorb* locus, it was also unexpected that the *Rorb^h1^* mutation can cause a pronounced gait disturbance while having almost no effect on transcription in the brain. This may suggest that *Rorb*-expressing inhibitory spinal interneurons are particularly sensitive to changes in *Rorb* abundance(Koch *et al*. 2017).

*Rorb*, along with other circadian-related genes, is thought to be involved in certain forms of bipolar disorder, epilepsy, and autism spectrum disorder (Mansour *et al*. 2009a; Mansour *et al*. 2009b; McGrath *et al*. 2009; Patel *et al*. 2010; McCarthy *et al*. 2012; Lesca *et al*. 2013; Lai *et al*. 2015; Konrad *et al*. 2019; Sadleir *et al*. 2020; Satterstrom *et al*. 2020). Based on these reports, we examined our RNAseq data, specifically the subsets of genes whose expression changed in *Rorb^h4/h4^*, for enrichment of genes linked to each of these disorders. All comparisons using gene-disease association data from OMIM and DisGeNET indicated significant overlap between these sets. Many gene products with high degree centrality in the PPI network have both high disease-association evidence scores and fold change magnitudes. This is particularly true in the differentially expressed genes of *Rorb^h4/h4^* with negative fold-changes (see *Grin2b*, DisGeNET evidence index 0.875/1.00). Negative log2FC genes, when queried alone, were associated with Mammalian Phenotype Ontology terms related to behaviors, including aberrant emotion/affect (Holm-Bonferroni p = 8.46e-10), fear/anxiety (Holm-Bonferroni p = 2.34e-7), and social investigation (Holm-Bonferroni p = 0.005). These terms, found in gene expression data from mice, bear resemblance to some symptoms of ASD and bipolar disorder. The set of molecular interactions portrayed in these PPI networks are likely important during formation of synaptic circuits and may provide additional insight into the utility of anticonvulsants in bipolar disorder(Grunze 2007). The set of genes overlapping between *Rorb^h2/h2^* and *Rorb^h4/h4^* contained terms that may be related to the unfolded protein response: chaperone cofactor-dependent protein refolding (Holm-Bonferroni p=0.0021), ‘de novo’ protein folding (Holm-Bonferroni p=0.0039), attenuation phase (Holm-Bonferroni p=0.0025), and HSF1-dependent transactivation (Holm-Bonferroni p=0.0135). These terms may be associated with cellular stress arising from disruption of neural circuit formation. The surprising number of genes associated with epilepsy, ASD, and bipolar disorder amongst differentially expressed genes shared between Rorb^h2/h2^ and Rorb^h4/h4^ mutants indicates that this gene set may play an important role in neural development. Given the link between *RORB* and epilepsy, ASD, and bipolar disorder, these mutant mice and our data, in particular the RNAseq data showing the altered expression of genes that occurs after mutational insult to both *Rorb* isoforms, will be of use in future studies that probe the complex molecular interactions underlying these common disorders.

Our findings demonstrate the power of a phenotype-driven approach: the mutations underlying a given phenotype often have important functional consequences, some of which may be subtle. Studying an overt phenotype such as gait can provide an inroad to unexpected biomedically relevant lines of inquiry. This allelic series of *Rorb* mice presents both an intriguing and cautionary example. First, it was necessary to perform whole genome sequencing in order to identify the causative mutation in *Rorb^h1/h1^* mice; a whole-exome sequencing experiment was performed first and the finding of 2x coverage on a single exon did not reach significance or raise this locus to our attention. This problem would be exacerbated for cases in which no mapping data and/or strong candidate genes exist. Second, the *Rorb^h1/h1^* mouse shows that there can be minimal gene expression changes despite having pronounced phenotypes. *Rorb^h2/h2^* and *Rorb^h4/h4^* data illustrates how a more severe mutational event in the same gene can cause profoundly different transcriptomic shifts that are relevant to human disease, and that changes in disease-related interaction networks may be non-linear with respect to overt phenotypes. Our results also exemplify how subtle developmental misregulation, while challenging to identify, may nevertheless disrupt processes that are crucial to neural circuit formation.

The cellular mechanisms underlying the “high stepper” or “duck gait” phenotype by which we originally identified these mice have been elucidated by Koch and colleagues, but it is clear that these animals display other interesting phenotypes relevant to neurodevelopment(Koch *et al*. 2017). Our findings also supplement those of Clark et al., which demonstrate the role of RORB in cell fate specification in the somatosensory cortex(Clark *et al*. 2020). We expect that our *Rorb* mice exhibit a spectrum of disruption in Rorb function that may be useful for further study of the role of *Rorb* in fate determination and function in cells of the CNS where complete loss-of-function models may not completely reflect those *RORB* variants that segregate with epilepsy and other neurodevelopmental conditions. For this reason, further characterization of neural circuit development in the somatosensory cortex and spinal cord of these mutants may be useful to improve understanding of the human conditions. Further, clarifying the nature of the interactions between disease-associated protein products in our PPI network from *Rorb^h4/h4^* mice may help identify crucially misregulated genes in the aforementioned neurodevelopmental conditions. Understanding the shared features of these disorders may help better explain the utility of anticonvulsant medications in bipolar disorder(Joshi *et al*. 2019). We hope that publication of this allelic series will aid investigators who are positioned to address these important questions.

## Authors’ Contributions

GCM, ALDT, and RWB designed the experiment. ALDT performed experiments with assistance from JC (Von Frey testing), MM and GCM (retinal histology), and CH and GCM (identifying the causative mutation). JB performed differential expression analysis on RNAseq data with guidance from EC. GCM performed additional enrichment and network analysis. BH maintained mice and provided expert phenotypic consultation. LR coordinated the whole-genome sequencing and identified the *Rorb^h1^* duplication and design, data analysis, and writing for ChIA-PET. OZ performed husbandry, embryo harvest, NPC culture, crosslinking, data collection, and writing for ChIA-PET. HT performed ChIA-PET data analysis. ML and CYN performed ChIA-PET sample processing. GCM, ALDT, and RWB analyzed data and wrote the manuscript. All authors read and approved the final manuscript.

## Acknowledgments

We thank The Jackson Laboratory’s Scientific Services, including Histology and Genome Technologies, for their expert technical assistance (Chia-Lin Wei), for their expert technical assistance, JAX Creative (Jen Torrance) for mouse photo/videography, and Computational Sciences (Vivek Philip) for analysis of RNAseq data. We thank Elissa Chesler for expert technical assistance. We thank Laurent Bogdanik for assistance with analyzing nerve images. This work was supported by F32 EY022825 (ALDT), NIH OD021325 (LGR). G. Murray was supported by the University of Maine institutional training grant 1T32GM132006-01 from the National Institute of General Medical Sciences (PIs: L. Liaw and C. Henry).

---

**Spreadsheet S1: All differentially expressed genes from *Rorb^h1/h1^, Rorb^h2/h2^*, and *Rorb^h4/h4^* sequencing relative to wildtype controls.** Differentially expressed genes with FDR < 0.05 and p-value < 0.05 are color-coded based on direction of differential expression by sign of the log2 fold-change (red – positive, green – negative).

**Figure S1:**
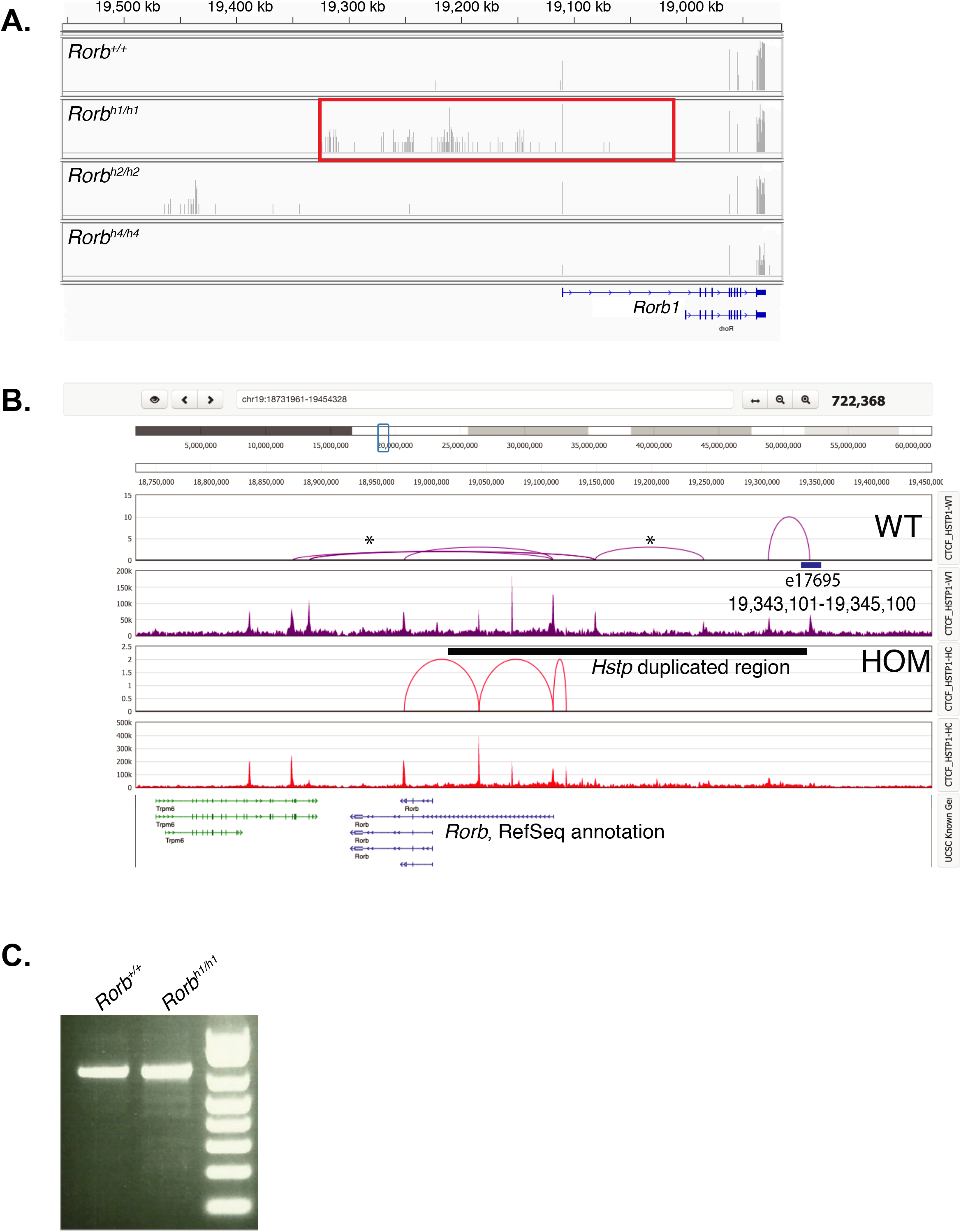
RNAseq and 5’RACE clarifies mutational events in *Rorb^h1/h1^* and *Rorb^h2/h2^* mice. Read depth in the region upstream of *Rorb* (A) reveals aberrant transcription in *Rorb^h1/h1^* mice, which is likely due to transcription occurring from the duplicated upstream *Rorb1* promoter and splicing into sequence within the duplicated region (red box). Similar aberrant transcription, albeit in a different pattern, is also seen in *Rorb^h2/h2^* mice. CTCF-mediated interactions in neural progenitor cells from *Rorb^hstp^/Rorb^hstp^ (HOM) and Rorb^+^/Rorb^+^ (WT) littermates*. CTCF-mediated interactions within and around the *Hstp* duplicated region are missing or rearranged in homozygous NPCs. The red peaks depict enrichment (normalized coverage) for DNA sequences bound by CTCF and the loops represent double-anchor factor bound supported chromatin loops. Shown in black is the duplicated *Hstp* region and in blue, a region containing strong distal enhancer (e17695) that is specific to developing forebrain (B). Despite this abnormal transcription pattern in *Rorb^h1/h1^* mice, 5’ RACE detected no altered transcripts in mutant brain (C). Animals were 6 weeks old, n=3 per genotype.

**Figure S2:**
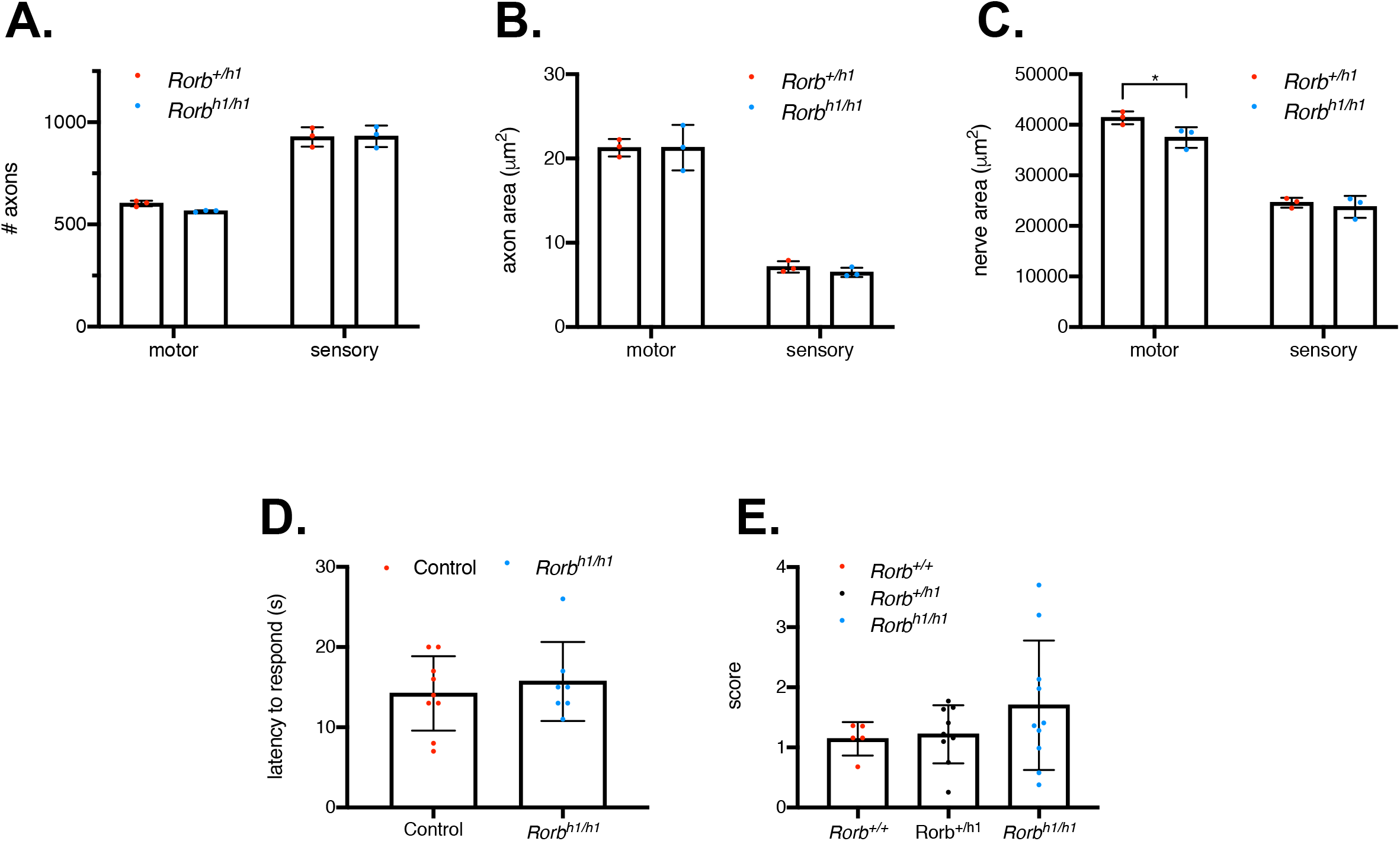
The *high stepper* phenotype is not associated with changes in femoral nerve anatomy or sensory behavior. Sensory and motor branches of the femoral nerve show no difference in axon count or diameter or overall nerve diameter (A-C) in *Rorb^h1/h1^* versus control mice. No differences between genotype were noted on the hot plate test of thermal nociception or the Von Frey test of mechanical sensation (D,E). Axon counts and average areas were obtained three 9-month-old mice per genotype. Hot plate: Animals were 8 months old, n=7 *Rorb^h1/h1^* and 9 littermate controls. Von Frey: Data were obtained from 2-month old animals, n=10 *Rorb^h1/h1^*, 9 *Rorb^+/h1^*, and 5 *Rorb^+/+^*. Data are shown ± SD, *p<0.05.

**Figure S3:**
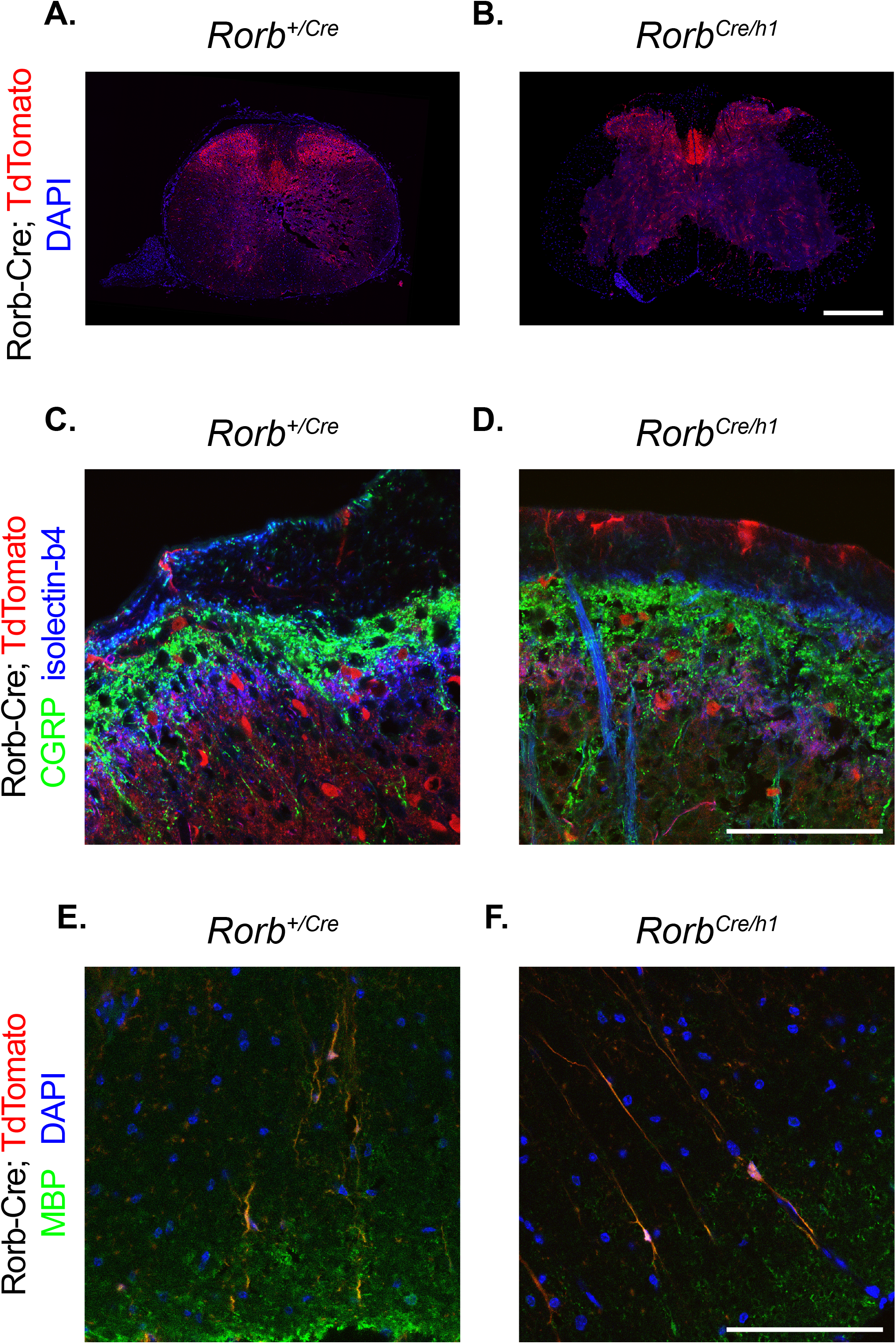
*Rorb* is expressed in regions of the dorsal horn associated with nociception and in oligodendrocytes. Sections of spinal cord from the cervical enlargement of mice carrying the *Rorb^Cre^; ROSA^tdT^* reporter system reveal prominent expression in the dorsal horn and in other scattered cells (A,B). The pattern and intensity of tdTomato signal was consistent between control (A) and *Rorb* mutant (B) mice. Co-imaging of tdTomato, CGRP, and isolectin b4 reveals tdTomato/isolectin b4 double-positive fibers in the dorsal horn in sections of spinal cord from the lumbar enlargement in both controls and mutants (C,D). Imaging for tdTomato and MBP reveals co-labeling within the white matter, indicating that the *Rorb^Cre^* reporter is expressed by at least some oligodendrocytes (E,F). Scale bar = 500 um in A-B, 100 um in C-F.

**Supplementary Table S1:** Pathway and process enrichment for *Rorb^h2/h2^*. All differentially expressed genes from *Rorb^h2/h2^* with FDR < 0.05 were uploaded for MouseMine gene list analysis. Reactome pathways, mammalian phenotype ontology, Gene Ontology biological process, and EMAPA anatomy enrichment terms with Holm-Bonferroni adjusted p-values and number of list matches are provided.

| Pathways | P-Value | Matches |
| --- | --- | --- |
| Collagen formation | 0.030813 | 6 |

| MP Term | P-Value | Matches |
| --- | --- | --- |
| Nervous system phenotype [MP:0003631] | 1.120859e-4 | 53 |
| Abnormal nervous system physiology [MP:0003633] | 0.006317 | 34 |
| Abnormal cell migration [MP:0003091] | 0.008827 | 14 |
| Abnormal nervous system morphology [MP:0003632] | 0.009635 | 42 |
| Abnormal bone structure [MP:0003795] | 0.014200 | 24 |
| Abnormal forebrain morphology [MP:0000783] | 0.030495 | 22 |

| GO Term | P-Value | Matches |
| --- | --- | --- |
| chaperone cofactor-dependent protein refolding | 0.036061 | 5 |

| EMAPA Term | P-Value | Matches |
| --- | --- | --- |
| molar | 6.724516e-4 | 25 |
| brain | 0.005612 | 103 |
| Nervous system | 0.020309 | 106 |
| Central nervous system | 0.026330 | 105 |
| cerebellum | 0.049412 | 61 |

*Rorb*<sup>h2/h2</sup> Anatomy (EMAPA) enrichment for all differentially expressed genes with FDR < 0.05. Holm-Bonferroni correction. Searched performed 5/9/2021.

**Supplementary Table S2:** Pathway and process enrichment for *Rorb^h4/h4^*. All differentially expressed genes from *Rorb^h4/h4^* with FDR < 0.05 were uploaded for MouseMine gene list analysis. Mammalian phenotype ontology, Gene Ontology biological process, Gene Ontology cellular process, Gene Ontology molecular function, and EMAPA anatomy enrichment terms with Holm-Bonferroni adjusted p-values and number of list matches are provided.

| MPO Term | P-Value | Matches |
| --- | --- | --- |
| abnormal emotion/affect behavior | 6.38E-08 | 67 |
| decreased body weight | 4.29678E-06 | 86 |
| decreased total tissue mass | 4.38801E-06 | 86 |
| abnormal behavior | 4.5416E-06 | 170 |
| behavior/neurological phenotype | 4.5416E-06 | 170 |
| abnormal glutamate-mediated receptor currents | 2.3988E-05 | 12 |
| decreased body size | 5.26829E-05 | 105 |
| abnormal body composition | 5.3725E-05 | 109 |
| abnormal body weight | 6.51434E-05 | 99 |
| abnormal postnatal growth/weight/body size | 0.000112977 | 137 |
| abnormal fear/anxiety-related behavior | 0.000147445 | 43 |
| abnormal body size | 0.000328853 | 119 |
| abnormal locomotor behavior | 0.000677812 | 91 |
| abnormal total tissue mass | 0.000825258 | 91 |
| abnormal nervous system physiology | 0.001915595 | 87 |
| abnormal voluntary movement | 0.004474886 | 93 |
| abnormal learning/memory/conditioning | 0.005423602 | 48 |
| abnormal anxiety-related response | 0.00553703 | 35 |
| abnormal cognition | 0.005685764 | 48 |
| abnormal NMDA-mediated synaptic currents | 0.008398522 | 7 |
| abnormal motor capabilities/coordination/movement | 0.008693224 | 123 |
| nervous system phenotype | 0.012775727 | 133 |
| hyperactivity | 0.015624077 | 43 |
| growth/size/body region phenotype | 0.029692883 | 182 |

| GO Term | P-Value | Matches |
| --- | --- | --- |
| anatomical structure morphogenesis | 2.12234E-05 | 113 |
| anatomical structure development | 4.93602E-05 | 192 |
| nervous system development | 7.85742E-05 | 93 |
| developmental process | 8.69651E-05 | 209 |
| multicellular organism development | 0.000318582 | 177 |
| positive regulation of cellular process | 0.000514277 | 193 |
| neuron projection development | 0.000518621 | 55 |
| system development | 0.000687708 | 153 |
| regulation of neuron projection development | 0.001308072 | 35 |
| positive regulation of biological process | 0.001667442 | 204 |
| cellular component organization or biogenesis | 0.00323805 | 207 |
| cellular component organization | 0.004184916 | 202 |
| neurogenesis | 0.006123426 | 67 |
| cell differentiation | 0.006265459 | 143 |
| regulation of cell development | 0.006708053 | 35 |
| plasma membrane bounded cell projection organization | 0.007694126 | 68 |
| cellular developmental process | 0.008603934 | 146 |
| positive regulation of metabolic process | 0.021581464 | 131 |
| cell projection organization | 0.024978938 | 68 |
| regulation of signaling | 0.030899473 | 121 |
| regulation of cellular component organization | 0.037656635 | 93 |
| regulation of cell communication | 0.044538191 | 120 |
| neuron projection morphogenesis | 0.044629085 | 36 |

| GO Term | P-Value | Matches |
| --- | --- | --- |
| postsynapse | 5.63417E-06 | 47 |
| organelle | 0.000122024 | 357 |
| intracellular anatomical structure | 0.000172397 | 388 |
| synapse | 0.00067041 | 68 |
| postsynaptic specialization | 0.002489928 | 29 |
| asymmetric synapse | 0.008343727 | 26 |
| neuron to neuron synapse | 0.009025084 | 27 |
| glutamatergic synapse | 0.009511773 | 30 |
| postsynaptic density | 0.011350804 | 26 |
| intracellular organelle | 0.016061991 | 329 |
| cell junction | 0.023747192 | 81 |
| postsynaptic membrane | 0.031544792 | 22 |
| dendrite | 0.038731677 | 37 |
| dendritic tree | 0.038731677 | 37 |
| extracellular matrix | 0.044418069 | 29 |
| external encapsulating structure | 0.047607211 | 29 |

| GO Term | P-Value | Matches |
| --- | --- | --- |
| protein binding | 3.39E-07 | 293 |
| binding | 0.023963785 | 372 |

| EMAPA Term | P-Value | Matches |
| --- | --- | --- |
| diencephalon | 3.25E-10 | 225 |
| hindbrain | 1.62E-09 | 253 |
| metencephalon lateral wall | 1.14E-08 | 219 |
| hypothalamus | 1.88E-08 | 165 |
| cerebellum | 2.00E-08 | 199 |
| metencephalon alar plate | 2.46E-08 | 200 |
| telencephalon | 5.37E-08 | 264 |
| thalamus | 1.07E-07 | 158 |
| brain | 1.65E-07 | 337 |
| metencephalon | 1.69E-07 | 221 |

Rorb<sup>h4/h4</sup> Anatomy (EMAPA) enrichment for all differentially expressed genes with FDR < 0.05. Holm-Bonferroni correction. Ranked by p-value. Top 10 terms of 70 that reached significance reported here ranked by p-value. Searched performed 5/9/2021.

**Supplementary Table S3:** Pathway and process enrichment for *Rorb^h4/h4^* positive and negative log2FC genes. Differentially expressed genes from *Rorb^h4/h4^* with FDR < 0.05 and negative or positive log2FC were uploaded separately for MouseMine gene list analysis. Reactome pathways and mammalian phenotype ontology with Holm-Bonferroni adjusted p-values and number of list matches are provided.

| Pathway | P-Value | Matches |
| --- | --- | --- |
| Neuronal System | 0.005794737 | 15 |
| LGI-ADAM interactions | 0.020258711 | 4 |

| MPO Term | P-Value | Matches |
| --- | --- | --- |
| abnormal emotion/affect behavior | 8.46E-10 | 44 |
| abnormal fear/anxiety-related behavior | 2.34E-07 | 29 |
| abnormal anxiety-related response | 5.40E-07 | 26 |
| decreased anxiety-related response | 0.00025514 | 15 |
| abnormal behavior | 0.000778766 | 77 |
| behavior/neurological phenotype | 0.000778766 | 83 |
| abnormal synaptic transmission | 0.001483937 | 28 |
| abnormal locomotor behavior | 0.004061634 | 45 |
| abnormal voluntary movement | 0.004427314 | 47 |
| abnormal social investigation | 0.00540249 | 10 |
| abnormal glutamate-mediated receptor currents | 0.005802286 | 7 |
| abnormal nervous system physiology | 0.009817167 | 43 |
| abnormal neuron morphology | 0.020505217 | 35 |
| abnormal synaptic plasticity | 0.022153168 | 7 |
| abnormal locomotor activation | 0.026567438 | 36 |
| hyperactivity | 0.045856114 | 23 |

| Pathway | P-Value | Matches |
| --- | --- | --- |
| Collagen biosynthesis and modifying enzymes | 0.013394104 | 8 |

| MPO Term | P-Value | Matches |
| --- | --- | --- |
| decreased body weight | 0.000218511 | 56 |
| decreased total tissue mass | 0.000221748 | 56 |
| preweaning lethality | 0.000510121 | 96 |
| mortality/aging | 0.001094485 | 117 |
| abnormal body weight | 0.001529645 | 64 |
| growth/size/body region phenotype | 0.003294168 | 122 |
| abnormal body composition | 0.003405741 | 69 |
| decreased body size | 0.005264075 | 66 |
| abnormal postnatal growth/weight/body size | 0.005782598 | 86 |
| abnormal total tissue mass | 0.009770296 | 59 |
| abnormal survival | 0.011714289 | 104 |
| abnormal body size | 0.012578521 | 75 |
| embryo phenotype | 0.021181355 | 51 |

Mammalian Phenotype Ontology enriched for *Rorb*<sup>h4/h4</sup> differentially expressed genes with a positive log2FC and FDR < 0.05 queried via MouseMine. Holm-Bonferroni correction. Searched performed 5/9/2021.

**Supplementary Video S1: High-stepper gait phenotype.** Video recording of a *Rorb^h5/h5^* mouse moving across a level surface. Hyperflexion of the rear limbs is evident.

## REFERENCES

B6;129S-Rorbtm1.1(cre)Hze/J, pp. Donating Investigator Hongku Zeng, Allen Institute for Brain Science. The Jackson Laboratory.

Differentially Expressed Gene Network Analysis, pp. Protocol. Cytoscape 3.8.2.

2020a Online Mendelian Inheritance in Man, OMIM®, pp. https://omim.org/. McKusick-Nathans Institute of Genetic Medicine, Johns Hopkins University (Baltimore, MD).

2020b RStudio Team, pp. RStudio, Boston, MA.

Amberger, J. S., C. A. Bocchini, F. Schiettecatte, A. F. Scott and A. Hamosh, 2015 OMIM.org: Online Mendelian Inheritance in Man (OMIM(R)), an online catalog of human genes and genetic disorders. Nucleic Acids Res 43:D789–798.

Andre, E., F. Conquet, M. Steinmayr, S. C. Stratton, V. Porciatti et al., 1998a Disruption of retinoid-related orphan receptor beta changes circadian behavior, causes retinal degeneration and leads to vacillans phenotype in mice. EMBO J 17:3867–3877.

Andre, E., K. Gawlas and M. Becker-Andre, 1998b A novel isoform of the orphan nuclear receptor RORbeta is specifically expressed in pineal gland and retina. Gene 216:277–283.

Baker, E. J., J. J. Jay, J. A. Bubier, M. A. Langston and E. J. Chesler, 2012 GeneWeaver: a web-based system for integrative functional genomics. Nucleic Acids Res 40:D1067–1076.

Becker, K. G., K. C. Barnes, T. J. Bright and S. A. Wang, 2004 The genetic association database. Nat Genet 36:431–432.

Bogdanik, L. P., J. N. Sleigh, C. Tian, M. E. Samuels, K. Bedard et al., 2013 Loss of the E3 ubiquitin ligase LRSAM1 sensitizes peripheral axons to degeneration in a mouse model of Charcot-Marie-Tooth disease. Dis Model Mech 6:780–792.

Bult, C. J., J. A. Blake, C. L. Smith, J. A. Kadin, J. E. Richardson et al., 2019 Mouse Genome Database (MGD) 2019. Nucleic Acids Res 47:D801–D806.

Byun, H., H. L. Lee, H. Liu, D. Forrest, A. Rudenko et al., 2019 Rorbeta regulates selective axon-target innervation in the mammalian midbrain. Development 146.

Carneiro, M., J. Vieillard, P. Andrade, S. Boucher, S. Afonso et al., 2021 A loss-of-function mutation in RORB disrupts saltatorial locomotion in rabbits. PLoS Genet 17:e1009429.

Chaplan, S. R., F. W. Bach, J. W. Pogrel, J. M. Chung and T. L. Yaksh, 1994 Quantitative assessment of tactile allodynia in the rat paw. J Neurosci Methods 53:55–63.

Clark, E. A., M. Rutlin, L. Capano, S. Aviles, J. R. Saadon et al., 2020 Cortical RORbeta is required for layer 4 transcriptional identity and barrel integrity. Elife 9.

Coppola, A., E. Cellini, H. Stamberger, E. Saarentaus, V. Cetica et al., 2019 Diagnostic implications of genetic copy number variation in epilepsy plus. Epilepsia 60:689–706.

Eppig, J. T., J. A. Blake, C. J. Bult, J. A. Kadin, J. E. Richardson et al., 2015 The Mouse Genome Database (MGD): facilitating mouse as a model for human biology and disease. Nucleic Acids Res 43:D726–736.

Fairfield, H., A. Srivastava, G. Ananda, R. Liu, M. Kircher et al., 2015 Exome sequencing reveals pathogenic mutations in 91 strains of mice with Mendelian disorders. Genome Res 25:948–957.

Feng, B., Y. Tang, B. Chen, C. Xu, Y. Wang et al., 2016 Transient increase of interleukin-1beta after prolonged febrile seizures promotes adult epileptogenesis through long-lasting upregulating endocannabinoid signaling. Sci Rep 6:21931.

Fu, Y., H. Liu, L. Ng, J. W. Kim, H. Hao et al., 2014 Feedback induction of a photoreceptor-specific isoform of retinoid-related orphan nuclear receptor beta by the rod transcription factor NRL. J Biol Chem 289:32469–32480.

Gnirke, A., A. Melnikov, J. Maguire, P. Rogov, E. M. LeProust et al., 2009 Solution hybrid selection with ultra-long oligonucleotides for massively parallel targeted sequencing. Nat Biotechnol 27:182–189.

Gorkin, D. U., I. Barozzi, Y. Zhao, Y. Zhang, H. Huang et al., 2020 An atlas of dynamic chromatin landscapes in mouse fetal development. Nature 583:744–751.

Grunze, H., 2007 [Anticonvulsants in the treatment of bipolar disorder]. Neuropsychiatr 21:110–120.

Gu, W., A. Wevers, H. Schroder, K. H. Grzeschik, C. Derst et al., 2002 The LGI1 gene involved in lateral temporal lobe epilepsy belongs to a new subfamily of leucine-rich repeat proteins. FEBS Lett 519:71–76.

Harris, C. R., K. J. Millman, S. J. van der Walt, R. Gommers, P. Virtanen et al., 2020 Array programming with NumPy. Nature 585:357–362.

Harris, J. A., K. E. Hirokawa, S. A. Sorensen, H. Gu, M. Mills et al., 2014 Anatomical characterization of Cre driver mice for neural circuit mapping and manipulation. Front Neural Circuits 8:76.

Jabaudon, D., S. J. Shnider, D. J. Tischfield, M. J. Galazo and J. D. Macklis, 2012 RORbeta induces barrel-like neuronal clusters in the developing neocortex. Cereb Cortex 22:996–1006.

Jia, L., E. C. Oh, L. Ng, M. Srinivas, M. Brooks et al., 2009 Retinoid-related orphan nuclear receptor RORbeta is an early-acting factor in rod photoreceptor development. Proc Natl Acad Sci U S A 106:17534–17539.

Joshi, A., A. Bow and M. Agius, 2019 Pharmacological Therapies in Bipolar Disorder: a Review of Current Treatment Options. Psychiatr Danub 31:595–603.

Kanehisa, M., and S. Goto, 2000 KEGG: kyoto encyclopedia of genes and genomes. Nucleic Acids Res 28:27–30.

Kautzmann, M. A., D. S. Kim, M. P. Felder-Schmittbuhl and A. Swaroop, 2011 Combinatorial regulation of photoreceptor differentiation factor, neural retina leucine zipper gene NRL, revealed by in vivo promoter analysis. J Biol Chem 286:28247–28255.

Kegel, L., E. Aunin, D. Meijer and J. R. Bermingham, 2013 LGI proteins in the nervous system. ASN Neuro 5:167–181.

Kindregan, D., L. Gallagher and J. Gormley, 2015 Gait deviations in children with autism spectrum disorders: a review. Autism Res Treat 2015:741480.

Koch, S. C., M. G. Del Barrio, A. Dalet, G. Gatto, T. Gunther et al., 2017 RORbeta Spinal Interneurons Gate Sensory Transmission during Locomotion to Secure a Fluid Walking Gait. Neuron 96:1419–1431 e1415.

Kohler, S., S. C. Doelken, C. J. Mungall, S. Bauer, H. V. Firth et al., 2014 The Human Phenotype Ontology project: linking molecular biology and disease through phenotype data. Nucleic Acids Res 42:D966–974.

Konrad, E. D. H., N. Nardini, A. Caliebe, I. Nagel, D. Young et al., 2019 CTCF variants in 39 individuals with a variable neurodevelopmental disorder broaden the mutational and clinical spectrum. Genet Med 21:2723–2733.

Lai, Y. C., C. F. Kao, M. L. Lu, H. C. Chen, P. Y. Chen et al., 2015 Investigation of associations between NR1D1, RORA and RORB genes and bipolar disorder. PLoS One 10:e0121245.

Lee, B., J. Wang, L. Cai, M. Kim, S. Namburi et al., 2020 ChIA-PIPE: A fully automated pipeline for comprehensive ChIA-PET data analysis and visualization. Sci Adv 6:eaay2078.

Lesca, G., G. Rudolf, N. Bruneau, N. Lozovaya, A. Labalme et al., 2013 GRIN2A mutations in acquired epileptic aphasia and related childhood focal epilepsies and encephalopathies with speech and language dysfunction. Nat Genet 45:1061–1066.

Li, H., and R. Durbin, 2009 Fast and accurate short read alignment with Burrows-Wheeler transform. Bioinformatics 25:1754–1760.

Lindskog, C., 2015 The potential clinical impact of the tissue-based map of the human proteome. Expert Rev Proteomics 12:213–215.

Liu, H., S. Y. Kim, Y. Fu, X. Wu, L. Ng et al., 2013a An isoform of retinoid-related orphan receptor beta directs differentiation of retinal amacrine and horizontal interneurons. Nat Commun 4:1813.

Liu, H., S. Y. Kim, Y. Fu, X. Wu, L. Ng et al., 2013b An isoform of retinoid-related orphan receptor beta directs differentiation of retinal amacrine and horizontal interneurons. Nature communications 4:1813.

Mansour, H. A., M. E. Talkowski, J. Wood, K. V. Chowdari, L. McClain et al., 2009a Association study of 21 circadian genes with bipolar I disorder, schizoaffective disorder, and schizophrenia. Bipolar Disord 11:701–710.

Mansour, H. a., M. E. Talkowski, J. Wood, K. V. Chowdari, L. McClain et al., 2009b Association study of 21 circadian genes with bipolar I disorder, schizoaffective disorder, and schizophrenia. Bipolar disorders 11:701–710.

McCarthy, M. J., C. M. Nievergelt, J. R. Kelsoe and D. K. Welsh, 2012 A survey of genomic studies supports association of circadian clock genes with bipolar disorder spectrum illnesses and lithium response. PLoS One 7:e32091.

McGrath, C. L., S. J. Glatt, P. Sklar, H. Le-Niculescu, R. Kuczenski et al., 2009 Evidence for genetic association of RORB with bipolar disorder. BMC Psychiatry 9:70.

Motenko, H., S. B. Neuhauser, M. O’Keefe and J. E. Richardson, 2015 MouseMine: a new data warehouse for MGI. Mamm Genome 26:325–330.

Patel, S. D., H. Le-Niculescu, D. L. Koller, S. D. Green, D. K. Lahiri et al., 2010 Coming to grips with complex disorders: genetic risk prediction in bipolar disorder using panels of genes identified through convergent functional genomics. American journal of medical genetics. Part B, Neuropsychiatric genetics : the official publication of the International Society of Psychiatric Genetics 153B:850–877.

Pinero, J., A. Bravo, N. Queralt-Rosinach, A. Gutierrez-Sacristan, J. Deu-Pons et al., 2017 DisGeNET: a comprehensive platform integrating information on human disease-associated genes and variants. Nucleic Acids Res 45:D833–D839.

Rouillard, A. D., G. W. Gundersen, N. F. Fernandez, Z. Wang, C. D. Monteiro et al., 2016 The harmonizome: a collection of processed datasets gathered to serve and mine knowledge about genes and proteins. Database (Oxford) 2016.

Rudolf, G., G. Lesca, M. M. Mehrjouy, A. Labalme, M. Salmi et al., 2016 Loss of function of the retinoid-related nuclear receptor (RORB) gene and epilepsy. Eur J Hum Genet 24:1761–1770.

Sadleir, L. G., G. de Valles-Ibanez, C. King, M. Coleman, S. Mossman et al., 2020 Inherited RORB pathogenic variants: Overlap of photosensitive genetic generalized and occipital lobe epilepsy. Epilepsia 61:e23–e29.

Satterstrom, F. K., J. A. Kosmicki, J. Wang, M. S. Breen, S. De Rubeis et al., 2020 Large-Scale Exome Sequencing Study Implicates Both Developmental and Functional Changes in the Neurobiology of Autism. Cell 180:568–584 e523.

Schaeren-Wiemers, N., E. Andre, J. P. Kapfhammer and M. Becker-Andre, 1997 The expression pattern of the orphan nuclear receptor RORbeta in the developing and adult rat nervous system suggests a role in the processing of sensory information and in circadian rhythm. Eur J Neurosci 9:2687–2701.

Srinivas, M., L. Ng, H. Liu, L. Jia and D. Forrest, 2006 Activation of the blue opsin gene in cone photoreceptor development by retinoid-related orphan receptor beta. Mol Endocrinol 20:1728–1741.

Staub, E., J. Perez-Tur, R. Siebert, C. Nobile, N. K. Moschonas et al., 2002 The novel EPTP repeat defines a superfamily of proteins implicated in epileptic disorders. Trends Biochem Sci 27:441–444.

Stelzer, G., N. Rosen, I. Plaschkes, S. Zimmerman, M. Twik et al., 2016 The GeneCards Suite: From Gene Data Mining to Disease Genome Sequence Analyses. Curr Protoc Bioinformatics 54: 1 30 31–31 30 33.

Szklarczyk, D., A. Franceschini, S. Wyder, K. Forslund, D. Heller et al., 2015 STRING v10: protein-protein interaction networks, integrated over the tree of life. Nucleic Acids Res 43:D447–452.

Tang, Z., O. J. Luo, X. Li, M. Zheng, J. J. Zhu et al., 2015 CTCF-Mediated Human 3D Genome Architecture Reveals Chromatin Topology for Transcription. Cell 163:1611–1627.

team, M. W. a. t. s. d., 2020a mwaskom/seaborn, pp. Zenodo.

Team, R. C., 2020b R: A language and environment for statistical computing, pp. https://www.R-project.org/. R Foundation for Statistical Computing, Vienna, Austria.

Uhlen, M., L. Fagerberg, B. M. Hallstrom, C. Lindskog, P. Oksvold et al., 2015 Proteomics. Tissue-based map of the human proteome. Science 347:1260419.

Wang, S., C. Sengel, M. M. Emerson and C. L. Cepko, 2014 A gene regulatory network controls the binary fate decision of rod and bipolar cells in the vertebrate retina. Dev Cell 30:513–527.

Welter, D., J. MacArthur, J. Morales, T. Burdett, P. Hall et al., 2014 The NHGRI GWAS Catalog, a curated resource of SNP-trait associations. Nucleic Acids Res 42:D1001–1006.

Yates, A. D., P. Achuthan, W. Akanni, J. Allen, J. Allen et al., 2020 Ensembl 2020. Nucleic Acids Res 48:D682–D688.

Zeilhofer, H. U., H. Wildner and G. E. Yevenes, 2012 Fast synaptic inhibition in spinal sensory processing and pain control. Physiol Rev 92:193–235.

Zhang, Y., C. H. Wong, R. Y. Birnbaum, G. Li, R. Favaro et al., 2013 Chromatin connectivity maps reveal dynamic promoter-enhancer long-range associations. Nature 504:306–310.

Zhou, G., O. Soufan, J. Ewald, R. E. W. Hancock, N. Basu et al., 2019 NetworkAnalyst 3.0: a visual analytics platform for comprehensive gene expression profiling and meta-analysis. Nucleic Acids Res 47:W234–W241.

